# Decreased Activity of the *Ghrhr* and *Gh* Promoters Causes Dominantly Inherited GH Deficiency

**DOI:** 10.1101/545384

**Authors:** Daisuke Ariyasu, Emika Kubo, Daisuke Higa, Shinsuke Shibata, Yutaka Takaoka, Michihiko Sugimoto, Kazunori Imaizumi, Tomonobu Hasegawa, Kimi Araki

## Abstract

Isolated growth hormone deficiency type II (IGHD2) is mainly caused by heterozygous splice-site mutations in intron 3 of the *GH1* gene. A dominant negative effect of the mutant growth hormone (GH) lacking exon 3 on wild-type GH secretion has been proposed; however, the molecular mechanisms involved are elusive. To uncover the molecular systems underlying GH deficiency in IGHD2, we established IGHD2 model mice, which carry both wild-type and mutant copies of the human *GH1* gene, replacing each of the endogenous mouse *Gh* loci. Our IGHD2 model mice exhibited growth retardation associated with intact cellular architecture and mildly activated ER stress in the pituitary gland, caused by decreases in the growth hormone releasing hormone receptor (*Ghrhr*) and *Gh* gene promoter activities. Decreases in *Ghrhr* and *Gh* promoter activities were likely caused by reduced levels of nuclear CREB3L2, which was demonstrated to stimulate the activity of the *Ghrhr* and *Gh* promoters. This is the first *in vivo* study revealing a novel molecular mechanism of GH deficiency in IGHD2, representing a new paradigm, differing from widely accepted models.

## Introduction

Isolated growth hormone deficiency type II (IGHD2) is a dominantly inherited growth hormone (GH) deficiency, first described in 1994, and mainly caused by heterozygous splice-site mutations in intron 3 of the *GH1* gene (Binder & Ranke, 1995; Cogan et al, 1994). The wild-type *GH1* allele transcript includes 5 exons and produces a 22-kDa wild-type GH protein; however, the mutant *GH1* allele transcript generates a 17.5-kDa exon 3 deletion-mutant GH (Δ3 GH), as a result of in-frame skipping of exon 3. The fact that patients harboring a deletion in one *GH1* allele exhibit normal stature indicates that a single wild-type *GH1* allele is sufficient to produce normal levels of wild-type GH secretion (Akinci et al, 1992); however, patients with IGHD2 have low serum concentrations of wild-type GH, despite having a wild-type *GH1* allele. Thus, it has been suggested that Δ3 GH exerts a dominant negative effect on wild-type GH secretion; however, the precise molecular mechanisms involved have remained elusive for more than 20 years.

Several *in vitro* studies have demonstrated that Δ3 GH is not secreted extracellularly (Graves et al, 2001; Iliev et al, 2005; Kannenberg et al, 2007; Mullis et al, 2002; Salemi et al, 2006), suggesting that the dominant negative effect of Δ3 GH is exerted within somatotropic cells of the pituitary, where GH is generated. Generally, mutant proteins exert dominant negative effects on secretory pathways of wild-type factors at the protein level (Deladoey et al, 2001; Ito et al, 1999; Jacobson et al, 1997), which has led many researchers to focus on wild-type and Δ3 GH protein interactions, such as heterodimer formation; however, no study has yet demonstrated definitive evidence of heterodimers comprising wild-type and Δ3 GH proteins. At present, it is widely accepted that Δ3 GH itself is not harmful to somatotroph (Graves et al, 2001), and that wild-type GH contributes to the degradation of Δ3 GH via protein interactions, leading to impairment of the wild-type GH secretory pathway (Kannenberg et al, 2007; McGuinness et al, 2003).

In contrast, previous *in vitro* studies revealed that Δ3 GH localizes to the endoplasmic reticulum (ER), due to its aberrant protein structure (Graves et al, 2001; Salemi et al, 2006), and is degraded by the proteasome (Ariyasu et al, 2013; Kannenberg et al, 2007), indicating that Δ3 GH potentially causes ER stress in the somatotroph. We previously demonstrated the involvement of ER stress and apoptosis in IGHD2, using rat GH4C1 cells stably expressing wild-type GH and Δ3 GH (Ariyasu et al, 2013), indicating that Δ3 GH itself impairs ER functions *in vitro*, without the involvement of wild-type GH, inconsistent with the widely accepted hypothesis described above.

Since GH secretion is regulated by growth hormone releasing hormone (GHRH) signaling, it is important to establish a usable model that includes the hypothalamus-pituitary axis. One *in vitro* study has authentically mimicked the hypothalamus-pituitary axis (Petkovic et al, 2010); however, *in vivo* animal models are imperative to clarify the molecular mechanisms involved in the GH deficiency of IGHD2. McGuinness *et al.* established Δ3 GH transgenic mice and reported that they exhibited widespread pituitary damage and severe macrophage invasion (McGuinness et al, 2003). This mouse model had been the sole *in vivo* model used to represent the human IGHD2 phenotype. *In vivo* studies have been performed using this model, including Δ3 GH knockdown by shRNA (Lochmatter et al, 2010), and altering splicing efficiency using butyrate (Miletta et al, 2016), which ameliorated impaired GH secretion; however, the mechanisms underlying IGHD2 have not been determined *in vivo* (Miletta et al, 2017).

To clarify the molecular processes causing impaired GH secretion in IGHD2, we established IGHD2 model mice by exchanging endogenous mouse *Gh* genes for the human wild-type *GH1 (wtGH1)* and mutant *GH1* (*Δ3GH1*; generates Δ3 GH) genes, using the ‘gene exchange system’ previously reported by our laboratory (Araki et al, 2002). Our IGHD2 model mice demonstrated significant growth failure, associated with a marked decrease in *wtGH1* mRNA, and no apoptosis was detected in the pituitary glands, despite the clear growth retardation phenotype.

Here, we show that Δ3 GH decreases transcription from the growth hormone releasing hormone receptor (*Ghrhr*) gene promoter, which has a fundamental role in somatotroph proliferation, as well as *GH1* gene transcription, leading to impaired GH production before birth in IGHD2 model mice. These decreases in promoter activity were mediated by a reduction in nuclear CREB3L2, one of the Creb3 family of bZip transcription factors, which was found to stimulate transcription from the *Ghrhr* and the *Gh* promoters coordinately with POU class 1 homeobox 1 (POU1F1). This is the first *in vivo* study to reveal a novel molecular mechanism underlying GH deficiency in IGHD2, and provides a new paradigm, different from the widely accepted model.

## Results

### IGHD2 model mice exhibited mild growth retardation

To establish a mouse model that authentically demonstrates the Δ3 GH-mediated dominant negative effect observed in IGHD2, one each of the mouse endogenous *Gh* gene alleles was exchanged for the human *wtGH1* and *Δ3GH1* genes, using the gene exchange system previously reported by our laboratory (Araki et al, 2002). Briefly, we inserted a *neoR* gene cassette, flanked by a left-element mutated *loxJT15* and *loxP* site, at the *Gh* gene locus by homologous recombination, producing a mouse *Gh* knock-out (KO) allele (*Gh^-^*) (Fig EV1A and C). Then, we constructed gene exchange vectors, containing right-element mutated loxKR3 and the *wtGH1* or *Δ3GH1* genes, followed by a puromycin resistance gene and the loxP site (Fig EV1A and C). The *neoR* gene cassette was exchanged for the *wtGH1* or *Δ3GH1* genes using Cre-mediated recombination (Fig EV1A and B). Since recombination between loxJT15 and loxKR3 produced a *lox* site with both sides mutated, which is resistant to Cre-mediated excision, we were able to efficiently generate recombined embryonic stem (ES) cell clones (Fig EV1A) (Araki et al, 2002).

Schematic representations of the original mouse endogenous *Gh* allele (Gh+), KO allele (Gh^-^), and exchanged human *GH1* alleles (*Gh^wtGH1^* or *Gh^Δ3GH1^*) are presented in Fig 1A. By crossing *Gh^+/-^, Gh^+/wtGH1^*, and *Gh^+/Δ3GH1^* mice, we successfully established *Gh^wtGH1/wtGH1^* mice (human healthy control model), *Gh^wtGH1/-^* mice (*GH1* heterozygous deletion model), *Gh^wtGH1Δ3GH1^* mice (IGHD2 model), and *Gh^-/-^* mice (*GH1* homozygous deletion model). The body weights and body lengths of these model mice are shown in Fig 1B and C. *Gh^-/-^* mice demonstrated severe postnatal growth retardation, associated with serum insulin-like growth factor 1 (IGF-1) levels below the detection range, indicating that, as in humans, postnatal growth in mice is dependent on GH activity (Fig 1B-E). *Gh^wtGH1/wtGH1^* and *Gh^wtGH1/-^* mice showed longitudinal growth and serum IGF-1 levels comparable with those of *Gh^+/+^* mice, suggesting that the human wild-type GH molecule is capable of binding the mouse GH receptor and producing IGF-1, and that one exchanged human wild-type *GH1* allele is sufficient for IGF-1-mediated longitudinal growth in mice (Fig 1B-E). *Gh^wtGH1/Δ3GH1^* mice exhibited mild growth retardation, associated with significantly reduced serum IGF-1 values, which were intermediate between those of *Gh^wtGH^-^/wtGH1^* and *Gh^-/^* mice (Fig 1B-E). These data indicate that *Gh^wtGH1/Δ3GH1^* mice successfully demonstrate the dominant negative effect of Δ3 GH, and that the growth retardation of *Gh^wtGH1/Δ3GH1^* mice is caused by impaired GH activity.

**Figure 1.**
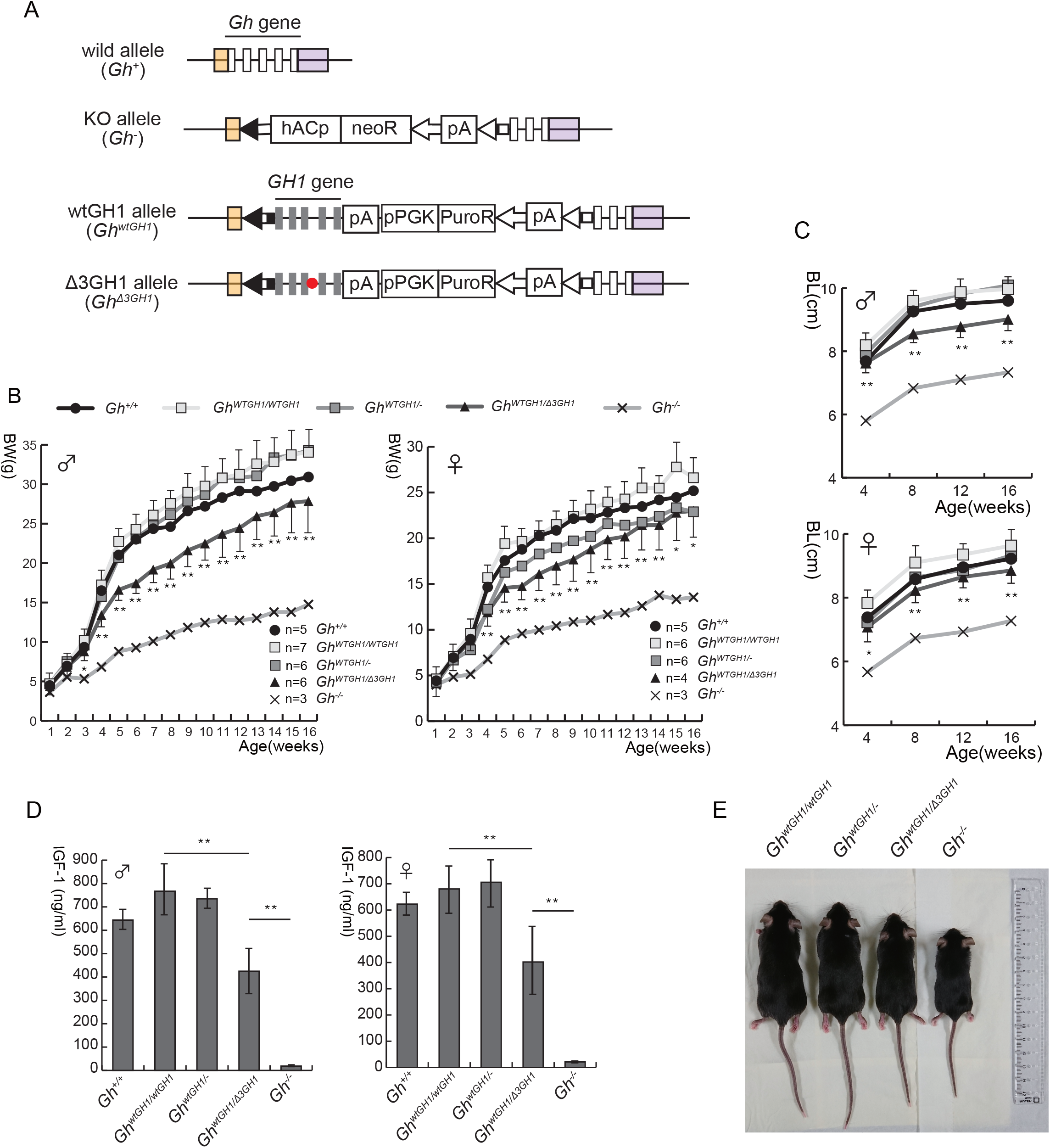
*Gh^wtGH1/Δ3GH1^* mice exhibit a Δ3 GH-mediated dominant negative phenotype. (A) Schematic representations of wild-type and genetically modified alleles. Vertical long open and closed rectangles represent the 5 exons of the endogenous *Gh* gene and the exchanged human *GH1* gene, respectively. Orange and purple rectangles, untranslated regions of the *Gh* gene. Red closed circle, c.291+1 g>a mutation. hACp, human actin promoter; neoR, neomycin resistance gene; pPGK, phosphoglycerate kinase promoter; PuroR, puromycin resistance gene; pA, polyadenylation signal. (B, C) Growth curves of model mice. (B) Body weight (BW) and (C) body length (BL) were measured every week and every 4 weeks, respectively, up to 16 weeks of age. Asterisks indicate that data from *Gh^wtGH1/Δ3GH1^* mice are significantly different to those from *Gh^wtGH1/wtGH1^* mice. Means calculated from the indicated numbers of model mice (bottom right in (B)) are shown. Error bars represent the standard deviation (SD) of values in *Gh^wtGH1/wtGH1^* and *Gh^wtGH1/Δ3GH1^* mice. *p<0.01, **p<0.005. (D) Serum IGF-1 values in 4-week-old model mice. Mean and SD values from six mice with each genotype are shown. **p<0.005. (E) Photograph of male *Gh^wtGH1/wtGH1^, Gh^wtGHy-^, Gh^wtGH1/Δ3GH1^*, and *Gh^-/-^* mice at 8 weeks old.

We also established *Gh^mGhΔ3GH1^* mice, in which the mouse *Gh* gene was inserted at the endogenous mouse *Gh* locus, using the gene exchange system in a similar way to that used to obtain *Gh^wtGH1Δ3GH1^* mice (Fig EV1A). *Gh^mGh/Δ3GH1^* mice also demonstrated growth failure, as for *Gh^wtGH1/Δ3GH1^* mice, although *Gh^+/Δ3GH1^* mice did not (Fig EV1E). The phenotypic discrepancy between *Gh+^Δ3GH1^* and *Gh^mGhΔ3GH1^* mice was attributable to a significant difference in the abundance of mRNA transcribed from the endogenous and exchanged *Gh* alleles, as demonstrated by qRT-PCR (Fig EV1F). In this study, we used mouse lines in which both alleles were exchanged for human *GH1* genes, because the transcriptional efficiencies of the *wtGH1* and *Δ3GH1* alleles were basically equivalent in human IGHD2 patients. These decreases in transcriptional efficiency of the exchanged human *GH1* genes did not cause the growth failure in *Gh^wtGH1/wtGH1^* mice (Fig 1B and C), and IGF-1 levels in *Gh^wtGH1/wtGH1^* mice were comparable with those in *Gh^+/+^* mice (Fig 1D), indicating that *wtGH1* mRNA expression levels were sufficient for production of human wild-type GH protein, required for the IGF-1-mediated longitudinal growth of mice.

### The growth retardation of *Gh^wtGH1/Δ3GH1^* mice is caused by decreased *wtGH1* mRNA expression

We evaluated the expression levels of wild-type and Δ3 GH proteins in pituitary glands at 4 weeks of age, the period in which *Gh^wtGH1/Δ3GH1^* mice demonstrated significant growth retardation, based on their growth curves (Fig 1B and C). Immunoblotting showed significant decreases in the content of the 22 kDa wild-type GH in whole *Gh^wtGH1/Δ3GH1^* pituitary, compared with *Gh^wtGH1/wtGH1^* and *Gh^wtGH1/-^* pituitaries (Fig 2A left and B), indicating that the growth retardation of *Gh^wtGH1/Δ3GH1^* mice was caused by impaired GH production in the somatotroph. Δ3 GH expression was barely detected by long exposure (Fig 2A right), despite comparable affinities of the anti-GH antibody for wild-type and Δ3 GH proteins (Fig EV2A). Immunostaining revealed that both the wild-type GH content in each somatotroph and the number of somatotrophs were reduced in *Gh^wtGH1/Δ3GH1^*, compared with *Gh^wtGH1/wtGH1^* pituitaries (Fig 2C).

**Figure 2.**
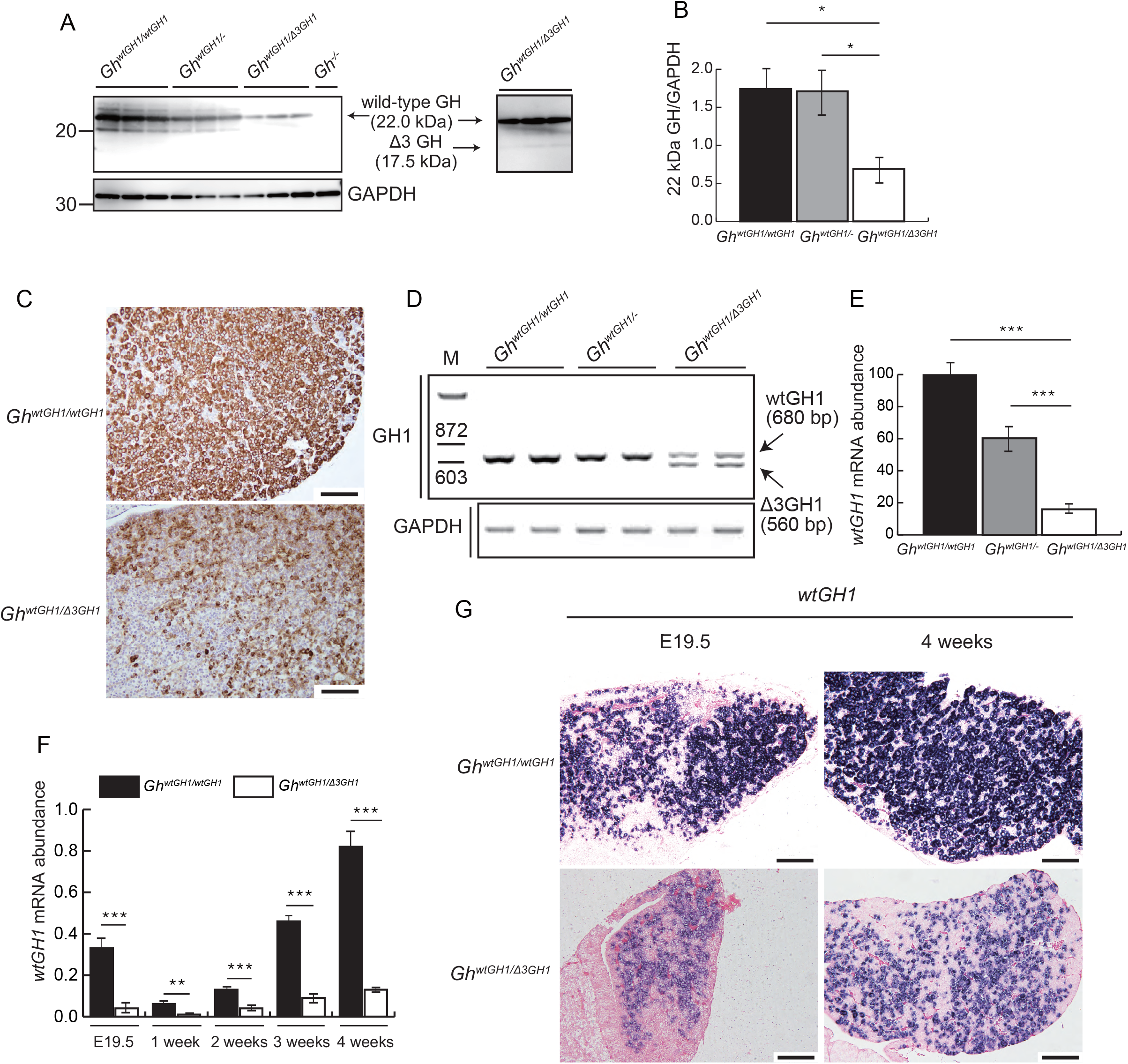
Evaluation of wild-type and Δ3 GH expression in *Gh^wtGH1/wtGH1^* and *Gh^wtGH1/Δ3GH1^* pituitary samples. (A) Immunoblotting using anti-GH antibody. This antibody recognizes Δ3 GH; however, the 17.5 kDa signals were too faint to be detected in *Gh^wtGH1/Δ3GH1^* pituitary glands. An image obtained by long exposure is shown on the right. (B) Evaluation of wild-type GH expression in (A) by densitometry. *p<0.05. (C) Results of immunohistochemistry using the same antibody used in (A). Scale bars, 100 μM. (D) Abundance of *wtGH1* and *Δ3GH1* mRNAs, evaluated by RT-PCR using a sense primer in exon 1 and an antisense primer in exon 5. (E) Evaluation of *wtGH1* mRNA abundance by qRT-PCR, using a sense primer in exon 3 and an antisense primer spanning exons 3 and 4. Data presented are mean and SD values from three independent samples, with relative abundance calculated by normalization to *β-actin* expression. ***p<0.005. (F) Results of qRT-PCR to evaluate the abundance of *wtGH1* mRNA using pituitary samples from *Gh^wtGH1/wtGH1^* and *Gh^wtGH1/Δ3GH1^* mice at E19.5, and 1, 2, 3, and 4 weeks of age. **p<0.01, ***p<0.005. (G) *In situ* hybridization of pituitary gland samples from E19.5 and 4-week-old mice, using an antisense probe complementary to *GH1* exon 3 mRNA. Scale bars, 100 μM.

Since we had demonstrated impaired production of wild-type GH in *Gh^wtGH1/Δ3GH1^* pituitary, we next evaluated *GH1* transcript levels. Using whole pituitary glands from 4-week-old animals, RT-PCR detecting both the *wtGH1* and *Δ3GH1* transcripts, with a sense primer in exon 1 and an antisense primer in exon 5, revealed markedly decreased *wtGH1* transcript levels in the *Gh^wtGH1/Δ3GH1^* pituitary, compared with those in *Gh^wtGH1/wtGH1^* and *Gh^wtGH1-^* pituitaries (Fig 2D). qRT-PCR analysis, detecting the *wtGH1* mRNA alone, using a sense primer in exon 3 and an antisense primer spanning exons 3 and 4, also demonstrated that the abundance of the *wtGH1* mRNA in *Gh^wtGH1/Δ3GH1^* pituitary was approximately one sixth of that in *Gh^wtGH1/wtGH1^* pituitary in 4-week-old animals (Fig 2E). Decreases in *wtGH1* mRNA levels in *Gh^wtGH1/Δ3GH1^* pituitaries were demonstrated from embryonic day E19.5 to 4 weeks of age (Fig 2F), suggesting that GH production in *Gh^wtGH1/Δ3GH1^* pituitaries was already impaired before birth. *In situ* hybridization analysis to detect *wtGH1* mRNA using an RNA probe for *GH1* exon 3, in E19.5 and 4-week-old pituitaries, revealed that both the abundance of *wtGH1* mRNA in each somatotroph and the number of somatotrophs were decreased in *Gh^wtGH1/Δ3GH1^*pituitaries, consistent with the results of immunostaining (Fig 2C and G). These data indicate that the impaired production of wild-type GH in *Gh^wtGH1/Δ3GH1^* pituitary is caused by a decrease in *wtGH1* mRNA.

RT-PCR showed that levels of the *wtGH1* and *Δ3GH1* transcripts were comparable (Fig 2D); however, immunoblotting revealed that expression of the Δ3 GH protein was drastically reduced compared with that of wild-type GH protein in *Gh^wtGH1/Δ3GH1^* pituitaries (Fig 2A). These data indicate that Δ3 GH is degraded in the somatotroph, and that wild-type GH is not involved in the degradation, inconsistent with the currently accepted hypothesis. Thus, to evaluate whether wild-type GH preferentially interacts with Δ3 GH, 3-D protein structures of wild-type and Δ3 GH were analyzed under pH conditions in the ER and docking simulation analysis was conducted using ZDOCK (Chen et al, 2003; Nakamura et al, 2017). The average binding affinity score for heterodimers of wild-type and Δ3 GH was significantly lower than that for the wild-type GH homodimer (Table 1), indicating that wild-type and Δ3 GH heterodimer formation is unlikely to be involved in impaired GH secretion in IGHD2.

**Table 1.** Results of docking simulation of wild-type and Δ3 GH. Higher ZDOCK score indicates more stable binding affinity. Numbers indicate times of the docking runs. ***p<0.001.

|  | wild type GH - wild type GH | wild type GH-Δ3 GH |
| --- | --- | --- |
| ZDOCK score | 150.6±16.58 (n=234) | 146.21±19.18 (n=938)* |

### Electron microscopy reveals markedly decreased numbers of secretory vesicles and enlarged ER in *Gh^wtGH1/Δ3GH1^* pituitaries

Considering the possibility that somatotroph loss due to apoptosis, necrosis, and inflammation, contribute to the decrease in *wtGH1* mRNA described above, we conducted histological evaluation of pituitary glands (Fig 3). Stereomicroscopic analysis of four-week-old *Gh^wtGH1/Δ3GH1^* pituitaries revealed a slightly atrophic and semi-translucent appearance, which was intermediate between those of *Gh^wtGH1/-^* and *Gh^-/-^* mice, consistent with the decreased somatotroph number in *Gh^wtGH1/Δ3GH1^* mice described above (Fig 3A); however, the decrease in somatotroph number was not caused by somatotroph loss, since hematoxylin-eosin staining demonstrated that the cellular architecture was intact, with no signs of necrosis or inflammation. Further, analysis by TdT-mediated dUTP nick end labeling (TUNEL) assay revealed that the decrease in the *wtGH1* mRNA in *Gh^wtGH1/Δ3GH1^* mice was not associated with apoptosis (Fig 3B).

**Figure 3.**
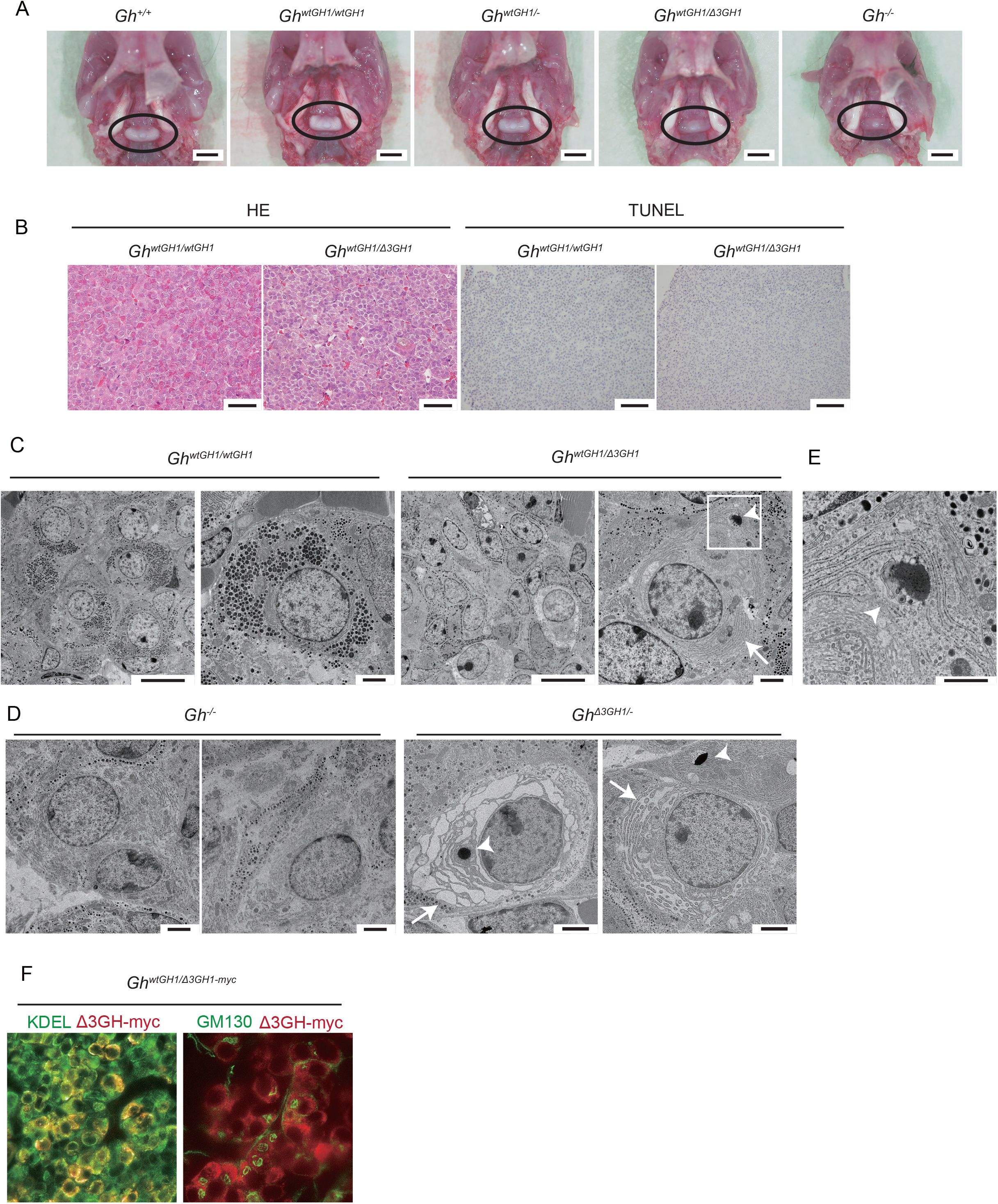
Histological evaluation of pituitary glands from *Gh^wtGH1/wtGH1^* and *Gh^wtGH1/Δ3GH1^* mice. (A) Stereomicroscope images of pituitary glands from 4-week-old *Gh^+/+^, Gh^wtGH1wtGH1^, Gh^wtGH1-^, Gh^wtGH1/Δ3GH1^*, and *Gh^-/-^* mice. Scale bars, 2 mm. (B) HE (scale bars, 50 μM) and TUNEL staining (scale bars, 100 μM) of pituitary glands from 4-week-old *Gh^wtGH1MGH1^* and *Gh^wtGH1/Δ3GH1^* mice. (C, D) TEM images of pituitary glands from 4-week-old (C) *Gh^wtGH1/wtGH1^* and *Gh^wtGH1/Δ3GH1^*, and (D) *Gh^-/-^* and *Gh^Δ3GH1/-^*, mice. Marked enlargement of the rough ER (arrow) and cytosolic protein aggregates (arrowhead) were observed in *Gh^wtGH1/Δ3GH1^* and *Gh^Δ3GH1/^* somatotroph. Scale bars, 10 μM (low power field) and 2 μM (high power field). (E) Magnified view of the open square in (C). The rough ER membrane is connected to the protein aggregates (arrowhead). Scale bar, 1 μM. (F) Evaluation of cellular localization of Δ3GH-myc by immunofluorescence, using antibodies against the myc-tag (Δ3GH-myc), KDEL (ER marker), and GM130 (Golgi marker).

The absence of somatotroph loss in *Gh^wtGH1/Δ3GH1^* mice led us to evaluate the characteristics of the cellular organelles in pituitary glands using transmission electron microscopy (TEM). TEM images of 4-week-old *Gh^wtGH1/wtGH1^* pituitary glands showed intact cell organelles, including appropriately developed rough ER, and many secretory vesicles containing mature wild-type GH protein (Fig 3C). In contrast, images of *Gh^wtGH1/Δ3GH1^* pituitary glands revealed a clear decrease in the number of secretory vesicles, abnormal enlargement of the rough ER, and protein aggregates in the cytosol (Fig 3C). To evaluate the impact of Δ3 GH itself on cellular morphology, we obtained *Gh^Δ3GH1/-^* mice by crossing *Gh^wtGH1/-^* and *Gh^wtGH1/Δ3GH1^* animals. In contrast to the current understanding that Δ3 GH itself is not harmful to somatotroph, TEM images of *Gh^Δ3GH1/-^* pituitaries revealed extreme enlargement of the rough ER and protein aggregates in the cytosol, whereas such abnormalities were not visible in *Gh^-/-^* pituitaries (Fig 3D). The observed protein aggregates were connected to the ER (Fig 3E), suggesting that the proteins accumulated in the ER were retro-translocated to the cytosol, leading to aggregate formation. These data led us to confirm the cellular localization of Δ3 GH by immunofluorescence analysis. Using *Gh^wtGH1/Δ3GH1-myc^* mice, expressing Δ3 GH with a C-terminal myc tag (Fig EV1A), Δ3 GH was also demonstrated to localize within the ER *in vivo* (Fig 3F).

To evaluate the contents of the protein aggregates in the cytosol, we dissociated *Gh^wtGH1/Δ3GH1^* anterior pituitary cells and separated them into soluble and insoluble fractions, in the presence or absence of treatment with the proteasome inhibitor, MG132, and evaluated the distributions of wild-type and Δ3 GH by immunoblotting, because several studies have demonstrated that cytosolic protein aggregates have low solubility (Ariyasu et al, 2013; Imai et al, 2001; Kannenberg et al, 2007; Ward et al, 1995). A significant proportion of Δ3 GH was detected in the insoluble fraction following MG132 treatment, although the majority of wild-type GH was sorted into the soluble fraction (Fig EV2B). These data suggest that ER-localized Δ3 GH is degraded by the proteasome, leading to Δ3 GH aggregation, which overwhelms the degradative capacity in the cytosol, and that most wild-type GH is not involved in the aggregates.

### Δ3 GH-mediated ER stress is not a direct cause of the growth failure of *Gh^wtGH1/Δ3GH1^* mice

The enlargement of the rough ER and Δ3 GH aggregates in the cytosol suggest that the *Gh^wtGH1/Δ3GH1^* somatotrophs are under ER stress, and we have previously shown that Δ3 GH causes ER stress to the somatotroph *in vitro* (Ariyasu et al, 2013). These data led us to investigate Δ3 GH-mediated ER stress *in vivo*. PKR-like endoplasmic reticulum kinase (PERK), activating transcription factor 6 (ATF6), and inositol requirement 1 (IRE1), are well-characterized ER membrane-located proteins which sense ER stress. In the presence of ER stress, PERK is activated by trans-autophosphorylation, ATF6 activates expression of the ER chaperone immunoglobulin heavy-chain binding protein (BiP), and IRE1 activates splicing of X-box binding protein 1 (*Xbp1*) mRNA via the PERK, ATF6, and IRE1 pathways, respectively (Ariyasu et al, 2017; Yoshida, 2007). PERK phosphorylation can be detected by immunoblotting, as phosphorylated PERK is associated with a mobility shift during SDS-PAGE (Harding et al, 1999). Immunoblotting revealed that the PERK was phosphorylated in 2 and 4-week-old *Gh^wtGH1/Δ3GH1^* pituitary glands (Fig 4A). Further, qRT-PCR demonstrated a significant increase in *BiP* mRNA abundance in 4-week-old *Gh^wtGH1/Δ3GH1^* pituitary (Fig 4B). *Xbp1* mRNA splicing was evaluated by competitive RT-PCR, using primers flanking the 26 bp sequence spliced out by IRE1α (Yoshida et al, 2001). *Xbp1* mRNA splicing was significantly increased at 1, 2, and 4 weeks of age in *Gh^wtGH1/Δ3GH1^* pituitaries (Fig 4C and D).

**Figure 4.**
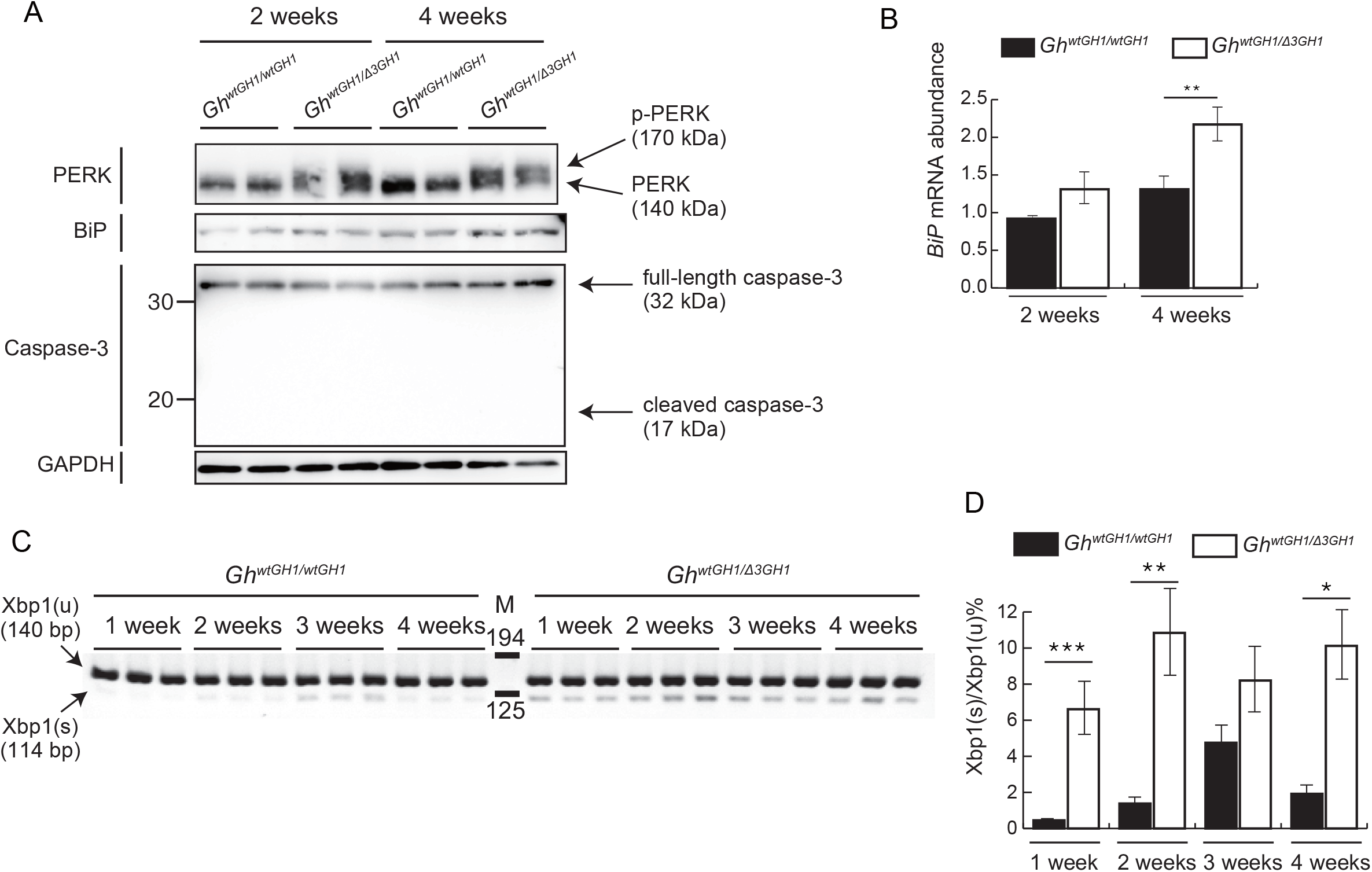
Δ3 GH activates three major ER stress response pathways. (A) Evaluation of phosphorylation of PERK, expression of BiP, and activation of caspase-3 by immunoblotting using pituitary glands from 2 and 4-week-old *Gh^wtGH1/wtGH1^*and *Gh^wtGH1/Δ3GH1^* mice. (B) qRT-PCR evaluation of the abundance of *BiP* mRNA in pituitary glands from 2 and 4-week-old *Gh^wtGH1/wtGH1^* and *Gh^wtGH1/Δ3GH1^* mice. **p<0.01. (C) RT-PCR evaluation of *Xbp1* mRNA splicing. Primers surrounding the region spliced out by IRE1 were used. PCR products of 140 and 114 bp, represent unspliced (*Xbp1*(*u*))and spliced (*Xbp1*(*s*)) *Xbp1*, respectively. M, PCR molecular weight marker. (D) Evaluation of the ratio of *Xbp1*(*s*) to *Xbp1*(*u*) by densitometry. *p<0.05, **p<0.01, ***p<0.005.

These data suggest that Δ3 GH can activate the three major ER stress pathways *in vivo;* however, activation of these ER stress pathways is not sufficiently strong to cause somatotroph apoptosis, since no apoptotic cells were detected by TUNEL assay, as described above (Fig 3B), and no caspase-3 activation was detected in *Gh^wtGH1/Δ3GH1^*pituitary glands by immunoblotting (Fig 4A). In other established ER stress-related endocrine diseases, apoptosis is required to cause organ dysfunction (Ariyasu et al, 2017; Fonseca et al, 2010; Hayashi et al, 2009; Oyadomari et al, 2002; Yoshida, 2007). These data suggest that Δ3 GH-mediated ER stress is not likely to be causally related to the growth retardation in *Gh^wtGH1/Δ3GH1^* mice.

### Decreased *Ghrhr* gene promoter activity contributes to decreased *wtGH1* gene expression in *Gh^wtGH1/Δ3GH1^* mice

Our data (described above) indicate that Δ3 GH itself can decrease *wtGH1* mRNA levels, without interacting with wild-type GH protein, ER stress, or apoptosis. RT-PCR revealed that both *wtGH1* and *Δ3GH1* mRNA were equally decreased in *Gh^wtGH1/Δ3GH1^* pituitary (Fig 2D), leading us to evaluate the expression levels of genes contributing to upstream regulation of *GH1* gene transcription.

As *wtGH1* mRNA was primarily decreased in *Gh^wtGH1/Δ3GH1^* somatotroph, we would expect the *Ghrhr* gene to be overexpressed, because of negative feedback mechanisms reflecting the GH deficiency in this tissue. Consistent with this hypothesis, 4-week-old *Gh^-/-^* mice had significantly increased abundance of *Ghrhr* mRNA compared with *Gh^wtGH1/wtGH1^* mice, because of a negative feedback mechanism, reflecting their complete GH deficiency (Fig 5A lane 1 and 3, and B). However, *Gh^wtGH1/Δ3GH1^* mice demonstrated significantly decreased *Ghrhr* mRNA compared with *Gh^wtGH1/wtGH1^* mice, despite their marked GH deficiency (Fig 5A lane 1 and 2, and B). These data suggest that the decrease in *wtGH1* mRNA in *Gh^wtGH1/Δ3GH1^* pituitary is mediated, at least in part, by that of *Ghrhr* mRNA. In agreement with this hypothesis, *Gh^Δ3GH1/-^* mice demonstrated significantly decreased abundance of *Ghrhr* mRNA, compared with *Gh^-/-^* mice, despite the fact that both strains lack the ability to secrete wild-type GH (Fig 5A lane 3 and 4, and B). Note that *Gh^Δ3GH1/-^* mice demonstrate exactly the same degree of growth retardation as *Gh^-/-^* mice, because Δ3 GH is not secreted (Fig 5C and EV2C). Further, the decrease in both the abundance of *wtGH1* mRNA in each somatotroph and the number of somatotrophs in *Gh^wtGH1/Δ3GH1^* pituitaries (Fig 2G), can be explained by this decrease in *Ghrhr* mRNA, because GHRH signaling is essential for *GH1* gene transcription and somatotroph proliferation (Lin et al, 1993). Taken together, Δ3 GH contributes to decreased *Ghrhr* mRNA levels, without the assistance of wild-type GH, leading to the impaired GHRH signaling and reduced *wtGH1* transcription in *Gh^wtGH1/Δ3GH1^* somatotroph.

**Figure 5.**
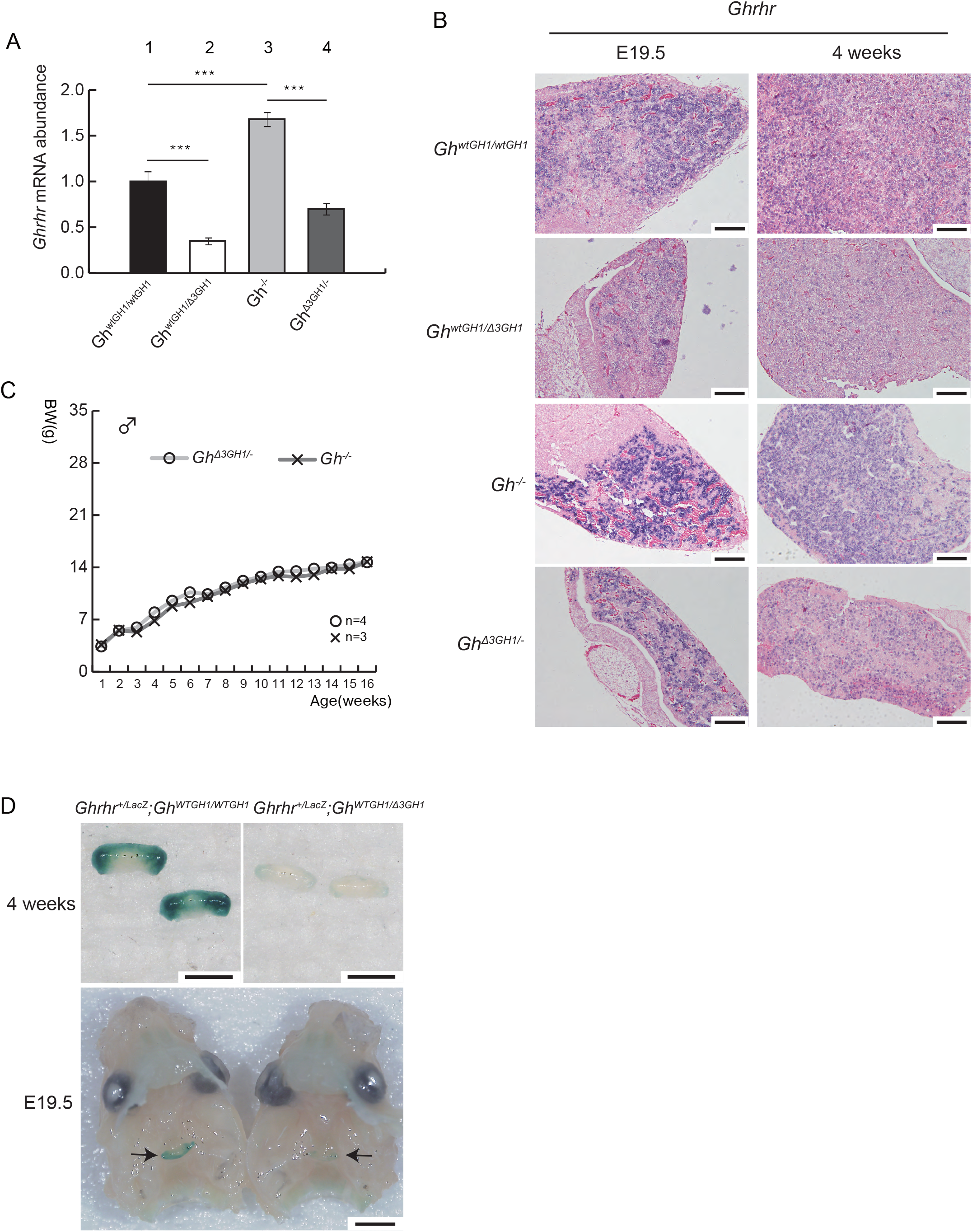
Δ3 GH decreases *Ghrhr* gene promoter activity. (A) Evaluation of *Ghrhr* mRNA abundance by qRT-PCR using samples from *Gh^wtGH1/wtGH1^*, *Gh^wtGH1/Δ3GH1^, Gh^-/-^*, and *Gh^Δ3GH1/-^* mice at 4 weeks old. Data presented are mean and SD values from three independent samples, with relative abundance calculated by normalization to *β-actin* expression. ***p<0.005. (B) *In situ* hybridization evaluation of *Ghrhr* mRNA expression in pituitary glands from E19.5 and 4-week-old *Gh^wtGH1/wtGH1^, Gh^wtGH1/Δ3GH1^*, *Gh^-/-^*, and *Gh^Δ3GH1/-^* mice. Scale bars, 100 μM. (C) Growth curves of male *Gh^-/-^* and *Gh^Δ3GH1/^* mice. (D) Results of X-gal staining of pituitary glands from 4-week-old and E19.5 *Ghrhr^+/LacZ^; Gh^wtGH1/wtGH1^* and *Ghrhr^+/LacZ^; Gh^wtGH1/Δ3GH1^* mice. Arrows indicate pituitary glands. Scale bars, 2 mm.

To evaluate the mechanisms underlying decreases in *Ghrhr* mRNA, a mouse model with the *LacZ* gene knocked in to the *Ghrhr* gene locus (*Ghrhr^+/LacZ^*) was established using the CRISPR/Cas9 gene editing system (Fig EV3A and B). E19.5 and 4-week-old *Ghrhr^+/LacZ^;Gh^wtGH1/Δ3GH1^* pituitary showed significantly decreased X-gal staining compared with *Ghrhr^+/LacZ^;Gh^wtGH1/wtGH1^* pituitary, indicating that the decreased *Ghrhr* mRNA levels are caused by a reduction in *Ghrhr* promoter activity (Fig 5D).

### Nuclear expressions of CREB3L2 is decreased in *Gh^wGH1/Δ3GH1^* pituitary glands

The decreased *Ghrhr* promoter activity detected in *Gh^wtGH1/Δ3GH1^* mice suggests that the expression of nuclear transcription factors crucial for the *Ghrhr* expression is disturbed by ER-localized Δ3 GH (Fig 5D). Furthermore, the abnormal cellular organelles and mildly activated ER stress pathway caused by Δ3 GH, indicate that Δ3 GH causes a deterioration in ER function by inducing ER stress (Fig 3C-E, Fig 4A-D). Immunoblotting revealed that abundance of the nuclear POU1F1 protein, a well-known transcription factor involved in regulation of *Ghrhr* and *Gh* promoter activities, was comparable in 4-week-old *Gh^wtGH1/wtGH1^* and *Gh^wtGH1/Δ3GH1^* pituitaries (Fig EV3C), suggesting that other, unknown, pituitary transcription factors were involved in the decreased *Ghrhr* promoter activity in *Gh^wtGH1/Δ3GH1^* mice. These data led us to focus on the Creb3 family of bZip transcription factors, a recently described family of ER stress transducers, all of which are ER-bound factors that undergo proteolysis in the Golgi apparatus, leading to production of active N-terminal fragments, which translocate to the nucleus and activate transcription of target genes (Kondo et al, 2011). Five Creb3 family members, CREB3/LUMAN, CREB3L1/OASIS, CREB3L2/BBF2H7, CREB3L3/CREBH, and CREB3L4/TISP40, have been identified to date (DenBoer et al, 2005; Kondo et al, 2005; Kondo et al, 2007; Nagamori et al, 2005; Omori et al, 2001).

Of these, CREB3L1 and CREB3L2 are essential for the differentiation and proliferation of osteoblasts and chondrocytes, respectively. Active N-terminal fragments of CREB3L1 and CREB3L2 bind to cAMP response element (CRE)-like sequences in the promoter regions of target genes, and enhance their expression in osteoblasts and chondrocytes, respectively (Murakami et al, 2009; Saito et al, 2009). Interestingly, *Creb3l1* KO mice display mild growth failure, which is not rescued by osteoblast-specific *Creb3l1* overexpression, but can be ameliorated by exogenous GH treatment, suggesting that they have impaired GH secretion (Murakami et al, 2011). Furthermore, TEM images of *Creb3l1*-deficient osteoblasts and Creb3l2-deficient chondrocytes showed abnormal enlarged ER (Murakami et al, 2009; Saito et al, 2009), similar to those detected in *Gh^wtGH1/Δ3GH1^* somatotroph. These similarities between *Gh^wtGH1/Δ3GH1^*and Creb3 family KO mice led us to hypothesize that one or more Creb3 family members are crucial for *Ghrhr* gene expression, and that the growth failure of *Gh^wtGH1/Δ3GH1^* mice is mediated by decreased levels of nuclear active N-terminal Creb3 protein fragments.

RT-PCR analysis of pituitary glands from 4-week-old C57BL/6 mice revealed that the *Creb3l1* and *Creb3l2* genes were strongly expressed in this tissue (Fig 6A). Next, we evaluated expression of these two genes in 4-week-old *Gh^wtGH1/wtGH1^* and *Gh^wtGH1/Δ3GH1^* pituitaries by qRT-PCR and immunoblotting. In *Gh^wtGH1/Δ3GH1^* pituitary, the abundance of *Creb3l1* mRNA was increased, while that of the N-terminal CREB3L1 protein was comparable with levels in the *Gh^wGH1wtGH1^* pituitary (Fig 6B and C). Furthermore, in *Gh^wtGH1/Δ3GH1^* pituitary, *Creb3l2* mRNA levels were comparable, while those of N-terminal CREB3L2 protein were reduced, compared with *Gh^wtGH1/wtGH1^* pituitary (Fig 6B and C). Both CREB3L1 and CREB3L2 were demonstrated to be expressed in somatotroph by immunofluorescence (Fig EV3D). These data indicate that expression levels of N-terminal CREB3L1 and CREB3L2 proteins are disturbed in *Gh^wtGH1/Δ3GH1^* mice.

**Figure 6.**
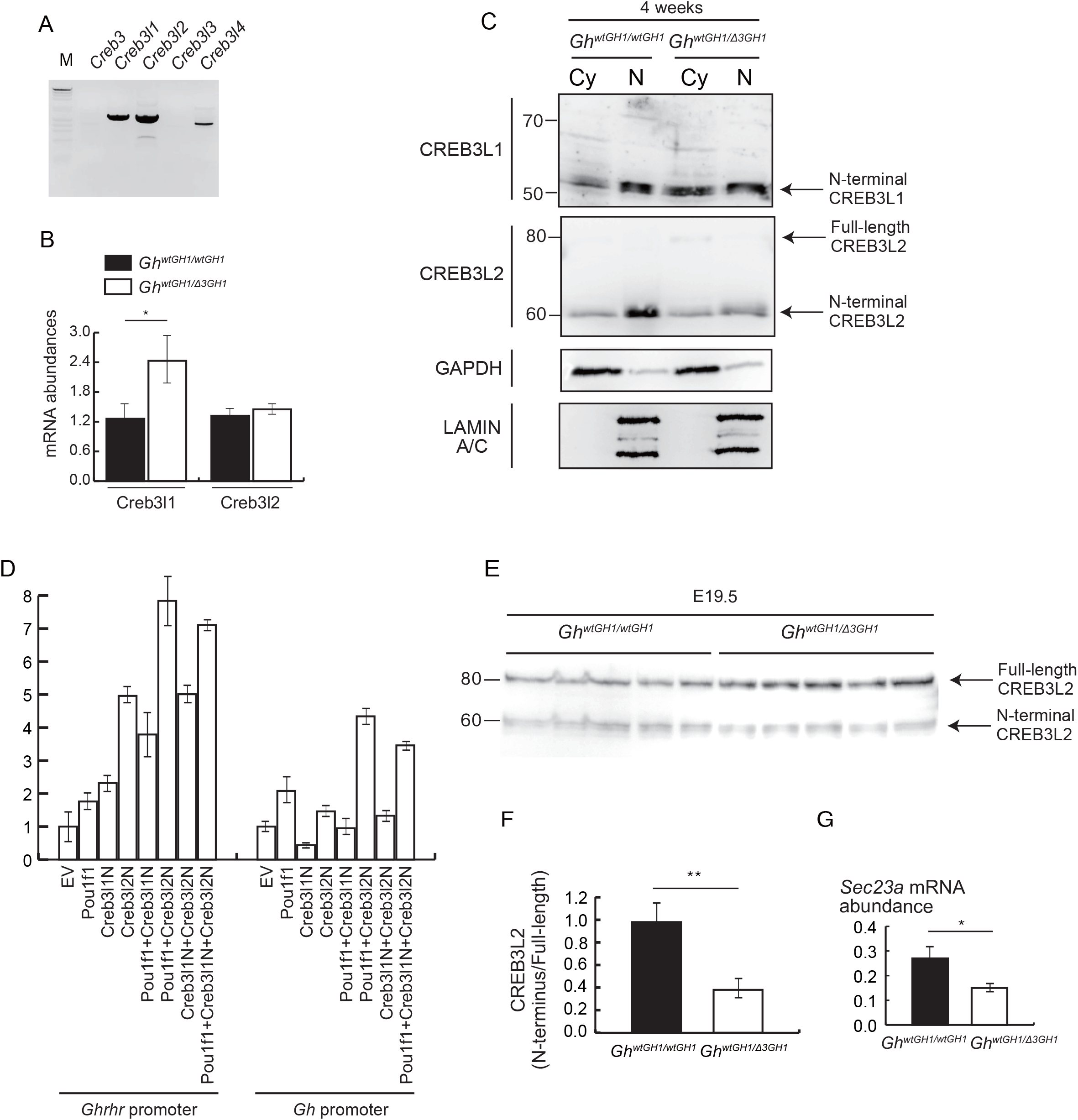
Levels of nuclear CREB3L2 are decreased in pituitary glands from *Gh^wtGH1/Δ3GH1^* mice. (A) Evaluation of Creb3 family member expression in 4-week-old whole pituitary glands by RT-PCR. (B) Evaluation of *Creb3l1* and *Creb3l2* gene expression in pituitary glands from 4-week-old *Gh^wtGH1/wtGH1^* and *Gh^wtGH1/Δ3GH1^* mice by qRT-PCR. *p<0.05. (C) Evaluation of cytoplasmic (Cy) and nuclear (N) expression levels of CREB3L1 and CREB3L2 by immunoblotting. Anterior pituitary cells from four-week-old *Gh^wtGH1/wtGH1^* and *Gh^wtGH1/Δ3GH1^* mice were dissociated and separated into Cy and N fractions. GAPDH and LAMIN A/C were used as positive controls for Cy and N fractions, respectively. (D) Reporter assay to evaluate expression driven by the *Ghrhr* (500 bp) and *Gh* (400 bp) promoters using vectors expressing POU1F1, CREB3l1-N, and CREB3l2-N. (E, F) Evaluation of CREB3L2 protein levels in pituitary glands from E19.5 mouse embryos by immunoblotting. **p<0.01. (G) Evaluation of the abundance of *Sec23a* mRNA in pituitary glands from E19.5 mouse embryos by qRT-PCR. *p<0.05.

Since involvement of N-terminal CREB3L1 and CREB3L2 in GH production has not been described previously, we evaluated the impact of N-terminal CREB3L1 and CREB3L2 on *Ghrhr* and the *Gh* promoter activities *in vitro*. Luciferase assays revealed that N-terminal CREB3L2 strongly stimulated transcription from both the *Ghrhr* and *Gh*promoters, in combination with POU1F1 (Fig 6D). Decreased nuclear CREB3L2 levels were also demonstrated at embryonic stages in *Gh^wtGH1/Δ3GH1^* pituitary glands by immunoblotting. Ratios of N-terminal CREB3L2 to full-length CREB3L2 decreased in *Gh^wtGH1/Δ3GH1^* pituitary glands at embryonic day E19.5 (Fig 6E and F), indicating that translocation of CREB3L2 to the nucleus was disturbed by ER-localized Δ3 GH. *Sec23a*mRNA, an established target gene of nuclear CREB3L2, was also decreased in *Gh^wtGH1/Δ3GH1^* pituitary at E19.5 (Fig 6G). These data suggest that CREB3L2 stimulates *Ghrhr* and the *Gh* gene expression levels, and that impaired GH production in *Gh^wtGH1/Δ3GH1^* somatotroph is mediated by a decrease in nuclear CREB3L2 levels.

## Discussion

The gene exchange system used in this study is a useful method for establishing mouse models of human diseases, because once researchers obtain a gene of interest flanked by mutated *lox* sites, they can easily exchange the endogenous genes with various human genes, via Cre-mediated integration. Using this system, we established a mouse model that expresses the *wtGH1* and *Δ3GH1* genes, instead of endogenous mouse *Gh*. *Gh^wtGH1/Δ3GH1^* mice have two advantages: 1) endogenous mouse GH is not produced and 2) one copy each of human wild-type GH and Δ3 GH are expressed under the control of the *Gh* promoter. Thus, *Gh^wtGH1/Δ3GH1^* mice are a genetically ideal model that mimics the growth patterns of human patients with IGHD2.

For functional analysis of secretory molecules regulated by feedback mechanisms involving in multiple tissues, such as GH, it is important to establish an applicable model that replicates the human disease as precisely as possible. Since patients with IGHD2 were first described in 1994, many researchers have focused on interference of wild-type GH protein trafficking by Δ3 GH, and several *in vitro* studies have been conducted to investigate the dominant negative effect of Δ3 GH (Ariyasu et al, 2013; Graves et al, 2001; Hayashi et al, 1999; Iliev et al, 2005; Kannenberg et al, 2007; Lee et al, 2000; McGuinness et al, 2003; Salemi et al, 2006; Salemi et al, 2007); however, in these studies, wild-type and Δ3 GH expression were driven by homeostatic promoters, such as the CMV promoter, and were thus independent of the feedback mechanisms that regulate GH expression *in vivo*. Furthermore, pituitary-derived cell lines, such as GC, GH3, GH4C1, and AtT-20, used in these studies do not express GHRHR. Thus, previous *in vitro* studies have been limited by the absence of feedback mechanisms mimicking those present *in vivo*.

The Δ3 GH transgenic mice reported by McGuinness demonstrated massive pituitary damage (McGuinness et al, 2003); however, our *Gh^wtGH1/Δ3GH1^* mice did not show similar somatotroph loss. The pituitary damage in the Δ3 GH transgenic mice is likely caused by overexpression of the transgene, compared with endogenous mouse *Gh*, since our data indicate that Δ3 GH caused GH deficiency, under conditions of comparable transcriptional efficiency between the mouse *Gh* and *Δ3GH1* alleles (Fig EV1E and F). Further, a previous study using cultured lymphocytes from patients with IGHD2 revealed that ratios of mutant to wild-type *GH1* transcripts were correlated with the severity of GH deficiency (Hamid et al, 2009).

In this study, we reveal two important findings: 1) Impaired GH secretion in *Gh^wtGH1/Δ3GH1^* mice is caused by decreased activity of the *Ghrhr* and *Gh* promoters; 2) these decreases in promoter activity are mediated by reduced levels of nuclear CREB3L2. A schematic representation of the molecular mechanisms underlying GH deficiency in IGHD2 is presented in Fig 7. Our data, including the abnormal TEM images and activated ER stress responses in *Gh^wtGH1/Δ3GH1^* pituitary, indicate that ER-localized Δ3 GH diminishes ER function by invoking ER stress. Δ3 GH-mediated ER stress leads to a decrease in COPII vesicles, which are essential for ER-Golgi transport, because Δ3 GH disturbs ER-Golgi transport (Graves et al, 2001) and COPII vesicles can be reduced under ER stress conditions (Shaheen, 2018). Full-length CREB3L2 is transported to the Golgi, where it is cleaved to produce the active N-terminal transcription factor (Kondo et al, 2011); thus a decrease in ER-Golgi transport is expected to reduce nuclear N-terminal CREB3L2 levels, leading to downregulation of both *Ghrhr* and *GH1* gene transcription. The decreased levels of N-terminal CREB3L2 disturb ER-Golgi transport in vicious cycle, since *Sec23a*, an established target gene of N-terminal CREB3L2, encodes SEC23, which is a COPII vesicle coat protein (Saito et al, 2009). Decreased ER-Golgi transport will also influence intracellular protein trafficking of wild-type GH and GHRHR, exacerbating the impaired secretion of wild-type GH. Furthermore, decreased GHRH signaling contributes to GH deficiency through inhibition of somatotroph proliferation. In contrast, decreased *Δ3GH1* gene transcription limits the accumulation of Δ3 GH in the ER, which likely protects the somatotroph from massive ER stress and apoptosis (Fig 7). To our knowledge, this is the first *in vivo* study that has come close to determining the molecular mechanisms underlying GH deficiency in IGHD2.

**Figure 7.**
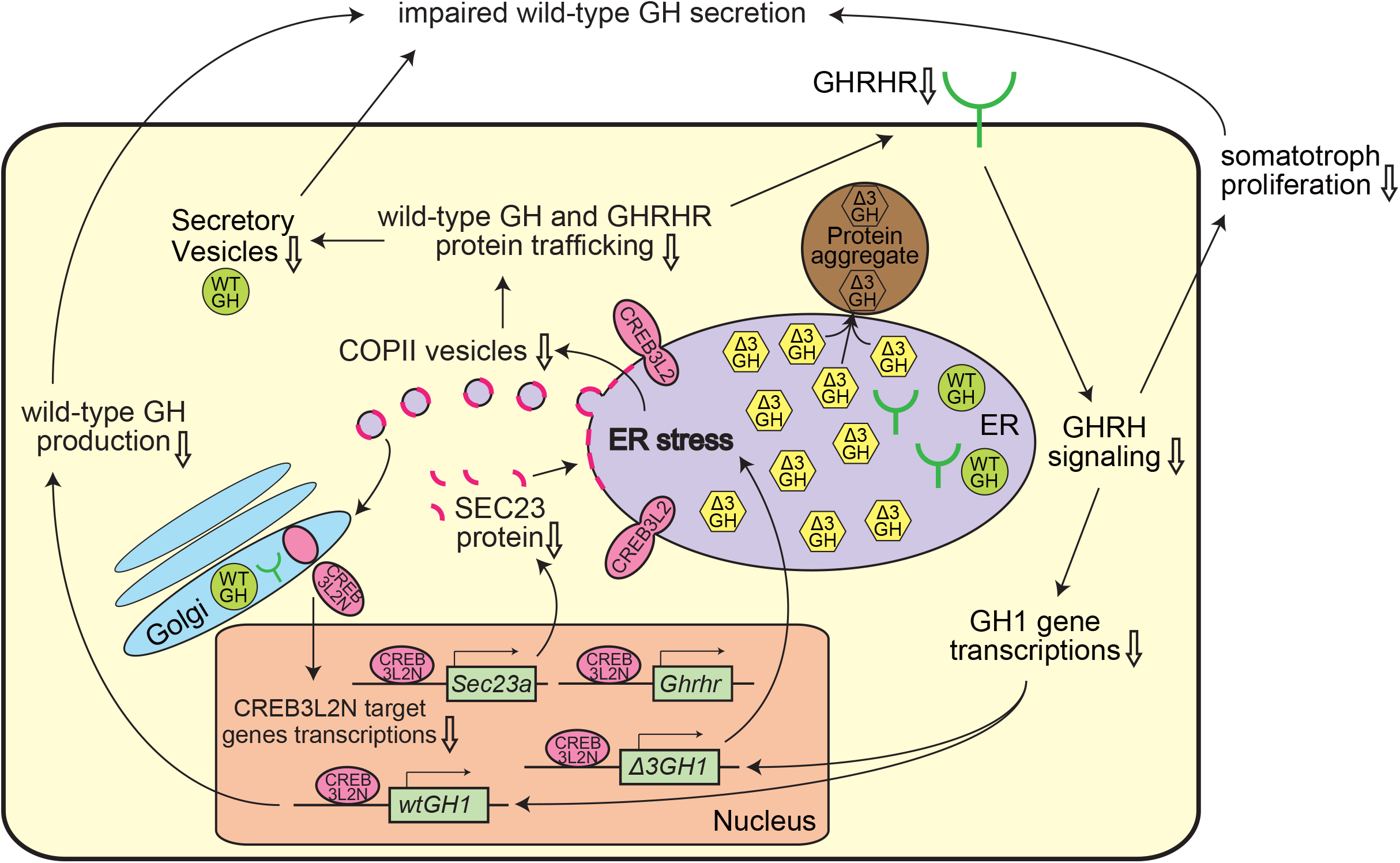
A schematic representation of Δ3 GH-mediated dominant negative effects in IGHD2 model mice.

The involvement of other transcriptions factors, such as CREB3L1 and/or unknown molecules, in the reduction of *Ghrhr* and *Gh* promoter activities is possible. Creb3 members can interact with other bZip transcription factors as heterodimers, to coordinately stimulate transcription of target genes (Saito et al, 2012; Vecchi et al, 2009; Zhang et al, 2006). Hence, identifying the partner molecule of CREB3L2 will be necessary for complete elucidation of the molecular mechanisms involved in IGHD2 GH deficiency.

We presume that CREB3L2 has an important role in the differentiation and proliferation of somatotroph at the late embryonic stage, by stimulating *GHRHR* and *GH1* gene transcription and increasing their secretion, similar to its role in chondrocytes. Postnatal growth has not been described in Creb3l2-deficient mice because systemic *Creb3l2* KO mice are lethal soon after birth (Saito et al, 2009). Thus, establishment of somatotroph-specific *Creb3l2* conditional KO mice, and/or *Gh^wtGH1/Δ3GH1^* mice overexpressing *Creb3l2* specifically in the somatotroph, would be warranted in the future.

Interestingly, exon 3 of the *GH1* gene has weak splice sites, and a small amount of Δ3 GH is produced in the pituitary glands, even in healthy individuals (Lecomte et al, 1987; Ryther et al, 2004). Δ3 GH may have a physiological role in preventing hyperproliferation of somatotroph and overexpression of the *GH1* gene, through the CREB3L2-mediated inhibition of *GHRHR* and *GH1* transcription.

This study has some limitations: 1) it remains unclear whether decreased activity of the *Ghrhr* and *Gh* promoters contributes to the GH deficiency in human IGHD2 patients, because patient pituitary samples are not available. Establishment of somatotrophs using induced pluripotent stem (iPS) cells derived from IGHD2 patients will help address this issue. 2) The observed decreases in transcriptional activity could be attributed to the influence of the gene exchange process (described in Fig EV1F) on amount of Δ3 GH protein produced, and the cellular characteristics of somatotrophs.

In this study, CREB3L1 and CREB3L2 were demonstrated to be involved in GH production in somatotrophs. These findings suggest that patients with GH deficiency may have mutations in the *Creb3l1* or *Creb3l2* genes, or their binding sites in the *Ghrhr* and *Gh* promoters.

In conclusion, IGHD2 model mice, created using our gene exchange system, reveal a novel molecular mechanism underlying GH deficiency, where ER-localized Δ3 GH leads to decreased levels of nuclear CREB3L2, and a consequent reduction in the activities of the *Ghrhr* and *Gh* promoters.

## Materials and Methods

### Isolation of the *wtGH1* and *Δ3GH1* genes

The *wtGH1* genomic sequence was amplified from healthy human control lymphocyte DNA using standard PCR methods with a sense primer (GH1-gf1) in the 5’-untranslated region (UTR) and an antisense primer (GH1-gr1) including the termination codon. The *Δ3GH1* genomic sequence, which contains a c.291+1 g>a mutation, was created by mutagenesis, using sense (GH1IVS3-F) and antisense (GH1IVS3dsmut-R) primers containing the single nucleotide substitution. None of the nucleotide polymorphisms, reported to be significantly associated with adult height (Hasegawa et al, 2000) were included in the sequences. All primers used in this study are listed in Supplementary Table 1.

### Plasmids

For KO of the *Gh* gene, the 5’ (7 kb) and 3’ (3 kb) arms were isolated by standard PCR methods, using a mouse bacterial artificial chromosome containing the *Gh* gene locus as a template, with sense (Gh-5arm-F and Gh-3arm-F) and antisense (Gh-5arm-R and Gh-3arm-R) primers. These 5’ and 3’ arms were inserted into the gene KO vector (Fig EV1A).

The *wtGH1, Δ3GH1, Δ3GH1-myc*, or *Gh* gene genomic sequences were inserted to the cassette exchange vector using appropriate restriction enzymes.

To establish the mouse model with *LacZ* knocked-in at the *Ghrhr* gene locus, 5’ (500 bp) and 3’ (450 bp) arms were isolated by standard PCR methods, using sense (Ghrhr-5arm-f5 and Ghrhr-3arm-f6) and antisense (Ghrhr-5arm-r5 and Ghrhr-3arm-r6) primers, and inserted into a plasmid containing the *LacZ* gene (Fig EV3A).

For CRISPR/Cas9 system plasmids, two pairs of oligonucleotides (Ghrhr-CRI-f1/r1, Ghrhr-CRI-f2/r2, Ghrhr-CRI-f4/r4, Ghrhr-CRI-f5/r5) were annealed and inserted into the pX335-U6-Chimeric_BB-CBh-hSpCas9n(D10A) plasmid (Addgene #42335) (pX335-Ghrhr-1, 2, 4, 5, and pX335-Rosa-3, 4).

For POU1F1, CREB3L1, and CREB3L2 expression plasmids their open reading frames (ORF) were isolated by standard PCR from C57BL/6 pituitary gland template cDNA using sense (Pou1f1-vf1, Creb3l1-vf1, and Creb3l2-vf1) and antisense (Pou1f1-vr1, Creb3l1-vr1, and Creb3l2-vr1) primers, and inserted into the pcDNA4 vector (Invitrogen).

For luciferase assay plasmids, the promoter regions of the *Ghrhr* (500 bp) and *Gh*(400 bp) genes were isolated from C57BL/6 mouse genomic DNA by standard PCR and inserted into the pGL4.10 vector (Promega).

For RNA probes to detect *GH1* and *Ghrhr* mRNAs by *in situ* hybridization, exon 3 of the *GH1* gene (120 bp) and the *Ghrhr* ORF (1272 bp) were amplified by PCR using sense (GH1ex3-f1 and Ghrhr-vf1) and antisense (GH1ex3-r1 and Ghrhr-vr1) primers, with C57BL/6 pituitary gland cDNA as a template, and inserted into the pSP73 vector (Promega).

### Embryonic stem cell culture and electroporation

KTPU8 feeder free ES cells, derived from F1 mice obtained by crossing C57BL/6 and CBA strains, were seeded into dishes pretreated with 0.15% gelatin solution, and cultured in Glasgow minimum essential medium, 14% knockout serum replacement, and 100 IU/ml leukemia inhibitory factor. To establish *Gh^+/-^* ES cells, KTPU8 cells cultured in a 10 cm culture dish were electroporated (0.8 kV/3 μF) with 20 μg linearized KO vector containing a neomycin resistance (*neoR*) gene flanked by the 5’ and 3’ arms, and seeded into four 10 cm culture dishes. Culture medium containing 180 μg/mL neomycin was exchanged daily. Nine days after electroporation, the surviving clones were picked and DNA from each clone was evaluated by Southern blotting (Fig EV1A and B).

To exchange the *neoR* gene for human *GH1* genes, the *Gh^+/-^* ES clones cultured in one 10 cm culture dish were electroporated (0.4 kV/125 μF) with 20 μg exchangeable vector and 10 μg Cre expression vector. Culture medium containing 0.8 μg/mL puromycin was exchanged daily, and the DNA samples of surviving clones were evaluated by Southern blotting (Fig EV1A and B).

To establish a mouse model in which the *LacZ* gene was knocked-in at the *Ghrhr*gene locus, ES cells derived from C57BL/6 mice were electroporated (0.4 kV/125 μF) with 25 μg plasmid containing the *LacZ* gene flanked by *Ghrhr* homology arms, pX335-Ghrhr-1 and pX335-Ghrhr-2 (15 μg each) (Fig EV3A). For this process, we took advantage of D10A mutant Cas9, to avoid making double-strand breaks (Cong et al, 2013; Jinek et al, 2012).

### Southern Blotting

Southern blotting was conducted using DIG, according to the manufacturer’s protocol. For *Gh* gene knockout, homologous recombination of the *neoR* gene was confirmed by Southern blotting using probes for sequences flanking the 5’ and 3’ arms, and the *neoR* gene (Fig EV1A and B). Substitution of the *GH1* gene for *neoR* was also evaluated by Southern blotting, using probes for the puromycin resistance gene (*PuroR*) (Fig EV1B). To evaluate knock-in of the *LacZ* gene at the *Ghrhr* gene locus, Southern blotting was performed, using probes for sequences flanking the 5’ and 3’ arms (Fig EV3A and B).

### RT-PCR and quantitative RT-PCR (qRT-PCR)

Total RNA was extracted from mouse pituitaries using an RNeasy mini kit (Qiagen) and 250 ng converted to cDNA using Revertra Ace (TOYOBO, Osaka, Japan, FSQ-101). cDNA aliquots were used for RT-PCR and qRT-PCR.

For qRT-PCR, THUNDERBIRD SYBR qPCR Mix (TOYOBO, QPS-201) was used as Taq polymerase and reactions were performed using the SYBR method on an Applied Biosystems 7500 Real-Time PCR System (Applied Biosystems, Foster City, CA). In experiments using relative quantification, the relative concentrations of target mRNAs were calculated using a standard curve and normalized to *β*-actin expression.

### SDS-PAGE and immunoblotting

SDS-PAGE and immunoblotting were carried out using polyacrylamide gels, polyvinylidene difluoride membranes, and ECL select detection reagent (Amersham, Buckinghamshire, England, RPN2235) according to standard procedures (Sambrook et al).

### Antibodies

For immunoblotting assays, the following primary antibodies were used: 1) anti-GH rabbit polyclonal antibody (Dako, A0570) (dilution 1:3000); 2) anti-PERK rabbit monoclonal antibody C33E10 (Cell Signaling Technology, 3192) (dilution 1:1000); 3) anti-caspase-3 rabbit monoclonal antibody 8G10 (Cell Signaling Technology, 9665) (dilution 1:1000); 4) anti-GRP78 rabbit monoclonal antibody EPR4041(2) (Abcam, ab108615) (dilution 1:500); 5) anti-GAPDH mouse monoclonal antibody 6C5 (Santa Cruz, sc-32233) (dilution 1:5000); 6) anti-LAMIN A/C rabbit polyclonal antibody (Cell Signaling Technology, 2032) (dilution 1:1000); 7) anti-POU1F1 mouse monoclonal antibody 2C11 (Abcam, ab10623) (dilution 1:1000); 8) anti-CREB3L1 mouse monoclonal antibody 44c7 (Merck Millipore, MABE1017) (dilution 1:1000); 9) anti-CREB3L2 rabbit polyclonal antibody (kindly provided by Prof. Imaizumi, Hiroshima University, Japan) (dilution 1:1000). Anti-rabbit immunoglobulin (IgG) (Dako, P0399) and anti-mouse immunoglobulin (Abcam, ab205719) (dilution 1:5000) secondary antibodies were used.

For immunohistochemistry, the following primary antibodies were used at a 1:100 dilution: 1) anti-GH rabbit polyclonal antibody (Dako, A0570); 2) anti-GH mouse monoclonal antibody (Abcam, ab15317); 3) anti-CREB3L1 rabbit polyclonal antibody (Abcam, ab33051); and 4) anti-CREB3L2 rabbit polyclonal antibody (kindly provided by Prof. Imaizumi). Goat anti-rabbit IgG (Abcam, ab150079, and Dako, P0448) (dilution 1:100) and M.O.M. biotinylated anti-mouse IgG reagent (VECTOR, MKB-2225) secondary antibodies were used. Fluorescein avidin DCS (VECTOR, A-2011) was used with the anti-mouse IgG reagent (MKB-2225).

### Luciferase assay

BMT10 cells were seeded into 96-well plates at a density of 5000 cells/well. Next day, plasmids expressing POU1F1, CREB3L1-N, and CREB3L2-N or empty vector were mixed in various combinations and transfected using Lipofectamine 2000 (Invitrogen). Forty-eight hours later, luciferase assay was performed using Dual-Luciferase Reporter Assay System (Promega).

### 3-D structural analyses of GH proteins

The 3-D structure of wild-type GH was obtained from the Protein Data Bank (PDB ID: 1AXI) and that of Δ3 GH was analyzed using the homology model function of MOE software (Chemical Computing Group Inc.), as described previously (Nakamura et al, 2017; Ogasawara et al, 2016). Hydrogen atoms were then added to each protein, using the protonate 3D function of MOE under ER pH conditions (Kim et al, 1998). These structures were then subjected to molecular mechanics (MM) calculations using MOE, with the AMBER99 force field, until the root mean square gradient was 0.01 kcal/mol/Å. After heating for 250 ps to attain 310 K as the starting temperature, a 5000 ps production run of the molecular dynamic (MD) simulation was performed at 310 K, with NPT ensemble using NAMD software (Phillips et al, 2005).

### Docking simulation analysis of dimeric GH molecules

Docking simulations were performed using ZDOCK (Chen et al, 2003) with residues within 10 Å of cysteine residues (numbers 79, 191, 208, and 215) defined as docking sites in each protein (Ogasawara et al, 2016; Yasui et al, 2013). Two thousand docking runs were performed for each pair of GH molecules and docking poses were classified according to the distance between cysteine residues in the two GHs. The number and binding affinities of docking poses which could form dimers via intermolecular disulfide bonds (Grigorian et al, 2005) were analyzed.

### Animals

ICR and C57BL/6 mice were purchased from CLEA Japan, Inc. Egg zona pellucida from E2.5 ICR mouse embryos were removed and aggregated with KTPU8 ES clones. The resulting blastocysts were transferred to the uteruses of ICR female mice mated with vasoligated male mice. F0 mice, with 100% chimerism, were crossed with C57BL/6 mice, to obtain N1 offspring. *Gh^wtGH1/wtGH1^* and *Gh^wtGH1/Δ3GH1^* mice were backcrossed to C57BL/6 mice for at least ten generations.

### Genotyping

To determine mouse genotypes, standard PCR reactions were performed using template DNA samples extracted from toe clips from 7-day-old mice (Fig EV1A and D). A reverse primer with the 3’ end converted to thymine from cytosine at the first base of the intron 3 in the *GH1* gene (primer No. 4, completely matching the *Δ3GH1* allele) was used. PCR reactions to detect both *wtGH1* and *Δ3GH1* alleles used an annealing temperature of 60°C, with those to detect only the *Δ3GH1* allele using an annealing temperature of 66°C (Fig EV1D). The sequences of all primers used in this study are listed in Table 1.

### Measurement of mouse body weight and length

Mouse body weights were measured every week from 1 to 16 weeks of age. Body lengths (distances from nose to anus) were measured under anesthesia, induced using isoflurane, every 4 weeks from 4 to 16 weeks of age, using a ruler.

### Measurement of mouse serum IGF-1 concentration

Blood samples were obtained from mouse orbital veins at 4 weeks of age, using a heparinized tube. Samples were centrifuged at 3000 rpm for 5 min, to collect serum samples collected. IGF-1 concentrations were evaluated using a Mouse/Rat IGF-1 ELISA Kit (ALPCO, 22-IG1MS-E01).

### Hematoxylin and eosin (HE) staining, TUNEL assay, and Immunohistochemistry

Pituitary glands were fixed in 4% paraformaldehyde (PFA) for 24 h at 15-25 °C, dehydrated through increasing concentrations of ethanol, equilibrated with xylene, embedded in paraffin wax, and sectioned at 4 μm. Pituitary sections were stained with hematoxylin and eosin and examined by light microscopy. TUNEL assays were performed using an ApopTag Peroxidase In Situ Apoptosis Detection Kit (Millipore, S7100). Thymus sections were used as positive controls. For immunostaining of GH, pituitary sections were deparaffinized, rehydrated, and treated with 20 μg/ml proteinase K as an antigen retrieval step. Antibodies are described in the ‘antibodies’ section above. Staining was performed using diaminobenzidine (DAB) and hematoxylin.

### Immunofluorescence assay

Pituitary glands were fixed in 4% PFA for 2 h at room temperature, dehydrated through increasing concentrations of sucrose, embedded in Optimal Cutting Temperature Compound, and sectioned at 6 μm. Frozen pituitary sections were subjected to immunofluorescence analysis.

### *In situ* hybridization

Pituitary glands were fixed in 4% PFA for 48 h at room temperature. Deparaffinization, cell conditioning, prehybridization, and stringency washing were automated using a staining workstation (Ventana Discovery XT). RNA probes were synthesized by *in vitro* transcription, and labelled with digoxigenin (DIG), using a DIG RNA labeling kit (Roche, 11175025910). Probes for *wtGH1* and *Ghrhr* mRNAs were manually hybridized at 0.5 ng/ml, 65°C for 6 h, and 10 ng/ml, 68°C for 6 h, respectively.

### Transmission electron microscopy

Pituitary glands from four-week-old mice were used for TEM observation, as described previously (Shibata et al, 2015). Briefly, tissues were dissected out and fixed in 2.5% glutaraldehyde in 0.1 M phosphate buffer (pH 7.4) for 24 h at 4°C. After 2 h post-fixation with 1% OsO_4_ and dehydration through ethanol, then acetone with n-butyl glycidyl ether (QY1) including a graded concentration of Epon with QY-1, they were embedded into 100% Epon. Following 72 h of polymerization in pure Epon, 70 nm ultrathin coronal pituitary gland sections were prepared on copper grids and stained with uranyl acetate and lead citrate for 10 min. Sections were observed under a TEM (JEOL JEM-1400 plus).

### X-gal staining of pituitary glands

Pituitary glands were removed from four-week-old mice and fixed in 4% PFA for 1 h on ice. Then, samples were permeabilized using rinse buffer (phosphate buffered saline (PBS) containing 2 mM MgCl_2_, 0.01% sodium deoxycholate, and 0.02% Nonidet P-40) for 2 h on ice, washed three times for 30 min in PBS, and stained in rinse buffer containing 5 mM potassium ferricyanide, 5 mM potassium ferrocyanide, and 1 mg/ml 5-Bromo-4-Chloro-3-Indolyl-β-D-Galactoside (X-gal) overnight at 37°C. For E19.5 embryo samples, heads were removed and fixed in 4% PFA for 30 min on ice. Then, pituitary glands were exposed by removing the skull bone and brain before permeabilization using rinse buffer.

### Dissociation of pituitary cells

Anterior pituitary cells were dissociated, according to published methods, using 0.5 % trypsin-EDTA (Oomizu et al, 1998).

### Soluble/insoluble fraction assay, cytoplasmic/nuclear fraction assay

For soluble/insoluble fraction assays, anterior pituitary cells were dissociated as described above, and 0.5 x 10^6^ cells seeded in DMEM with 10% FBS in a 24-well plate with 10 μM MG132 (Peptide Institute, Osaka, Japan), or DMSO, for 4 h. Detailed procedures have been described elsewhere (Ariyasu et al, 2013). To separate cytoplasmic and nuclear fractions, a NE-PER Nuclear and Cytoplasmic Reagent Kit (Thermo Fischer Scientific, 78833) was used.

### Statistics

Data were analyzed by 2-tailed unpaired Student’s *t* test for comparisons of two groups. For all bar graphs, means ± SD are plotted.

### Study approval

The Animal Care Committee and Institutional Biosafety Committee of the Kumamoto University approved all mouse protocols. All experiments were performed in accordance with the Declaration of Helsinki and were approved by the Kumamoto University Ethics Committee for Animal Experiments (authorization number in Kumamoto University: C23-262, C24-278).

## Acknowledgments

This work was supported by JSPS KAKENHI Grant Numbers JP16H06276 and JP16H07081. The authors thank Mayumi Muta, Kumiko Murakami, Mai Nakahara, and Riki Furuhata for assistance with embryo manipulation; Michiyo Nakata and Sayoko Fujimura for assistance with preparing paraffin-embedded and frozen pituitary samples; Hiderou Yoshida for giving important advice regarding ER stress-related experiments; Haruo Nogami for providing information about *Ghrhr* promoter activity; and Yukihiro Hasegawa for fruitful discussions about the manuscript and long-standing support.

## Author contributions

DA and KA conceived the study. DA and KA designed the experiments. DA, EK, DH, YT, and SS performed the experiments and analyzed the data. DA, SS, and YT wrote the initial draft of the paper. TH, MS, KI, and KA critically revised the manuscript. All authors reviewed and edited the manuscript.

## Expanded View Figure Legends

**Expanded View Figure 1.**
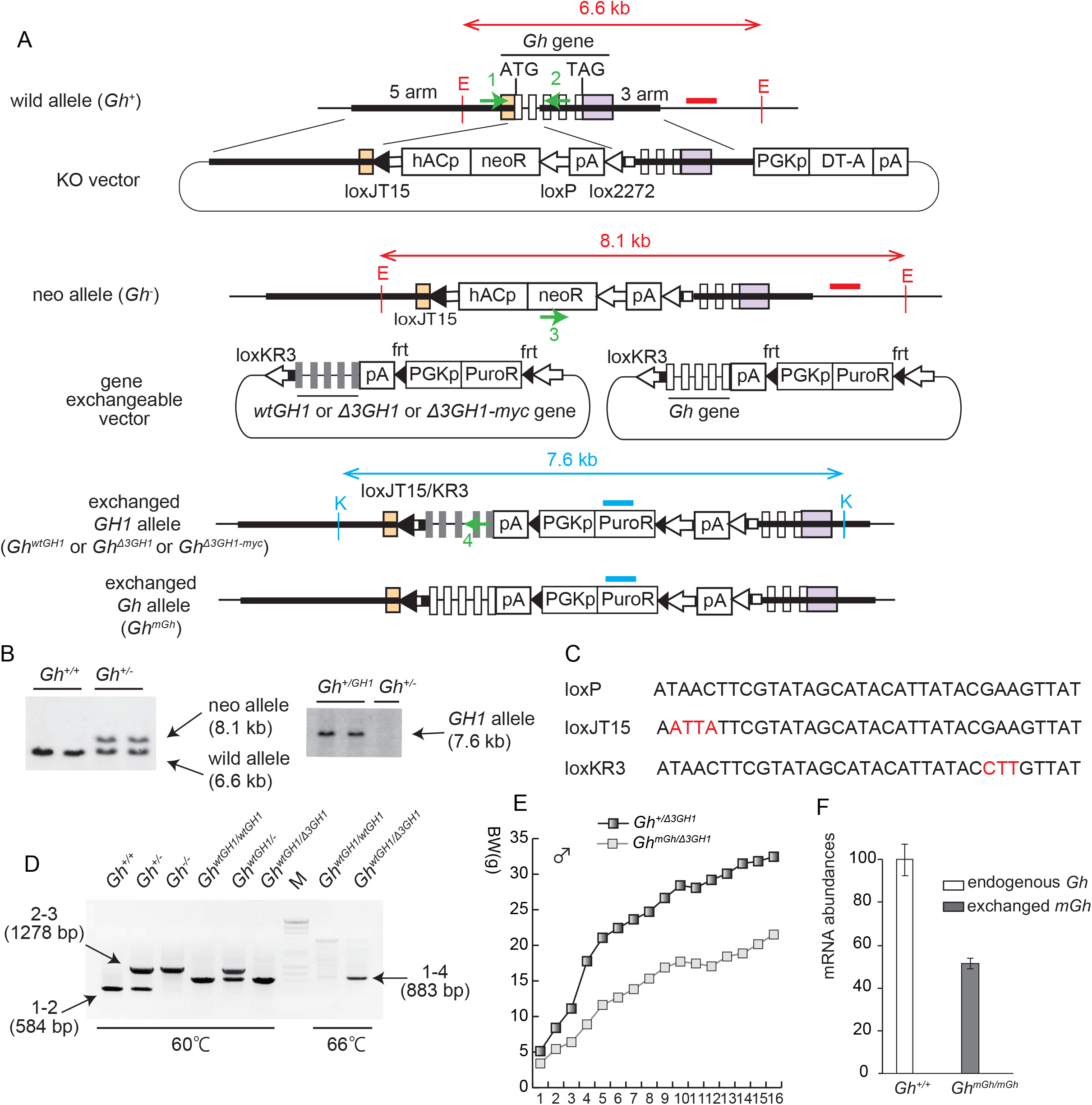
Establishment of IGHD2 model mice using the gene exchange system. (A) Schematic representation of the gene exchange system. Vertical long open and grey rectangles represent the open reading frames of the endogenous *Gh* and exchanged human *GH1* genes, respectively. Orange and purple rectangles represent untranslated regions of the *Gh* gene. Black bold lines represent homology arms. Red and blue bold lines represent probes used for Southern blotting to detect homologous recombination of the neoR gene and exchange of the *GH1* gene. Red and blue vertical lines represent EcoRI and KpnI recognition sites, respectively. Green arrows numbered from 1 to 4, genotyping primers. DT-A, diphtheria toxin A. (B) Results of Southern blotting using probes indicated in (A). (C) Intact and mutant loxP sequences used in this study; red letters indicate mutant sequences.(D) Typical results of PCR genotyping using primers 1 to 4. Signals detecting *Gh^+^, Gh^-^*, and *Gh^wtGH1^* or *Gh^Δ3GH1^* are shown. M: PCR molecular weight marker. (E) Growth curves of male *Gh+^Δ3GH1^* and *Gh^mGhΔ3GH1^* mice. (F) qRT-PCR evaluating the abundance of mRNA transcribed from the endogenous and exchanged *Gh* alleles in pituitary glands from 4-week-old mice.

**Expanded View Figure 2.**
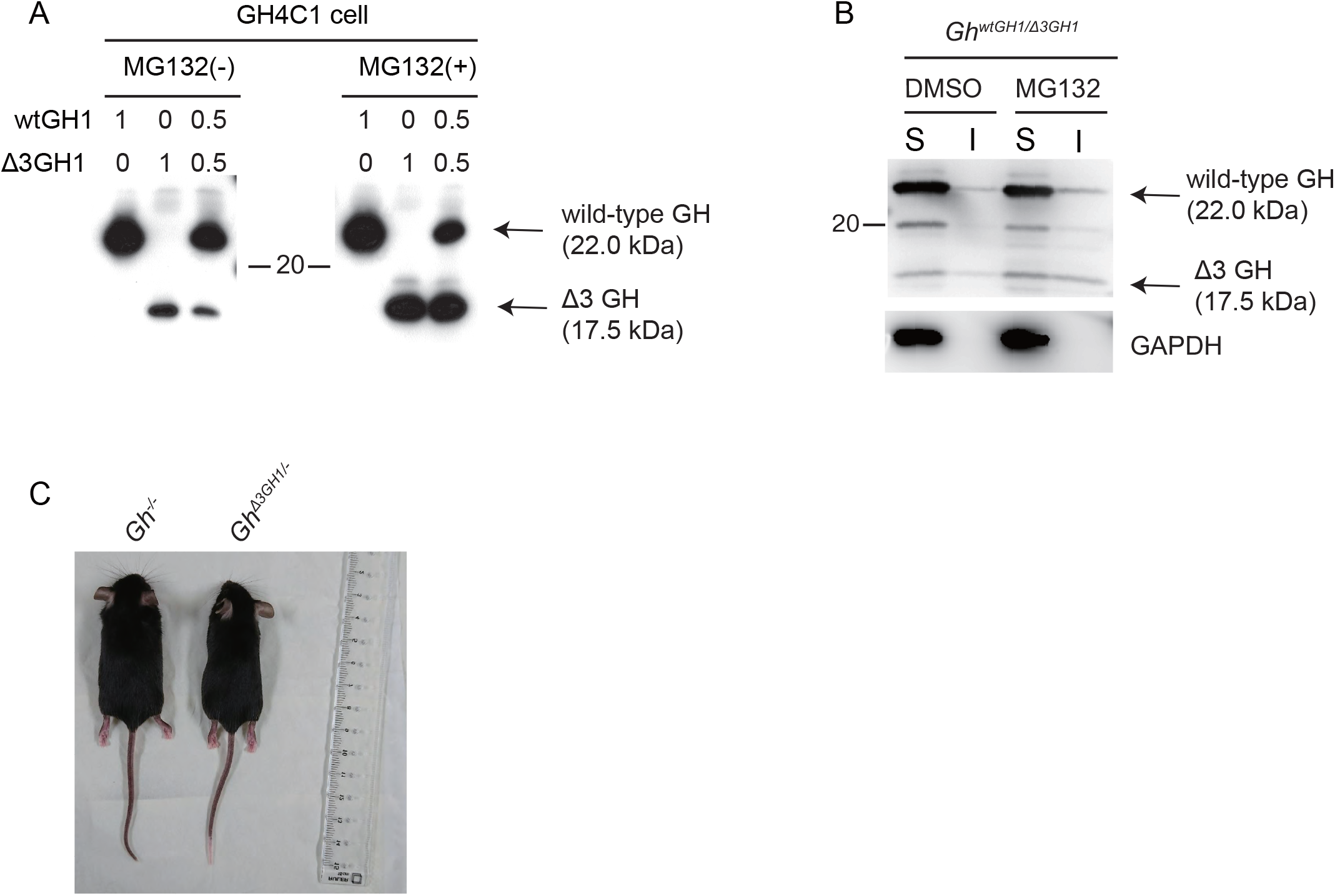
(A) GH4C1 cells were transfected with *wtGH1* and/or *Δ3GH1* cDNA constructs in the presence or absence of treatment with the proteasome inhibitor, MG132. Cell lysate samples were subjected to immunoblotting analysis using anti-GH antibody. Upper numbers indicate the quantities of transfected plasmid DNA, encoding either *wtGH1* or *Δ3GH1*, cDNA. Note that Δ3 GH is degraded by the proteasome and the anti-GH antibody used in this study has comparable affinities for both wild-type and Δ3 GH proteins. (B) Evaluation of the distribution of wild-type and Δ3 GH by soluble/insoluble fraction assay. Four-week-old anterior pituitary cells from *Gh^wtGH1/Δ3GH1^* mice were dissociated, separated into soluble (S) and insoluble (I) fractions, and subjected to immunoblotting in the presence or absence of MG132 treatment (10 μM, 4 h). GAPDH was used as a positive control for the soluble fractions. (C) Photograph of male *Gh^-/-^*, and *Gh^Δ3GH1/^* mice at 8 weeks old.

**Expanded View Figure 3.**
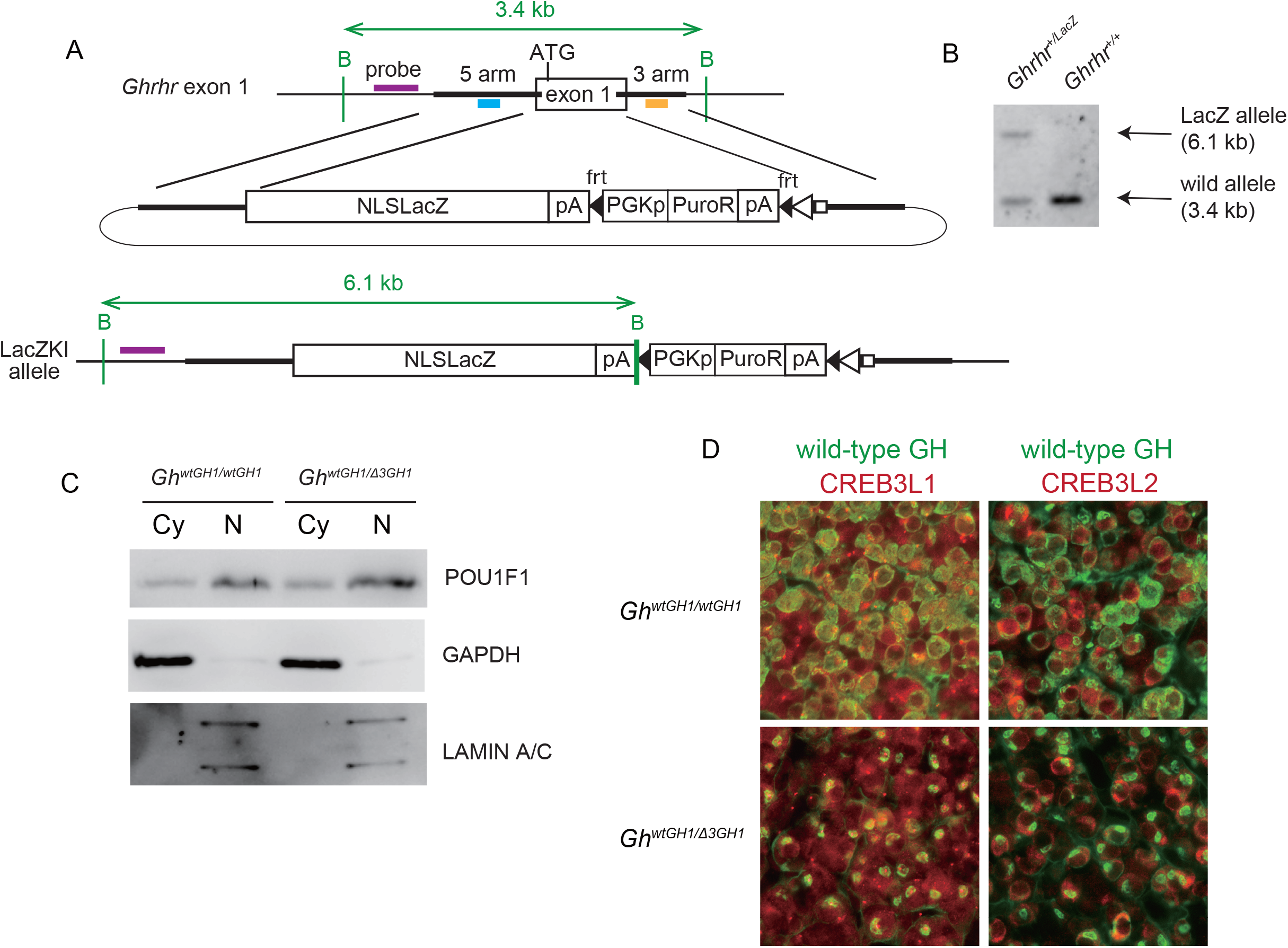
(A) A strategy for establishing *LacZ* knocked-in model mice, using the CRISPR/Cas9 system. The *LacZ* gene was inserted into exon 1 of the *Ghrhr* gene by homologous recombination. Blue and orange bars, 5’ and 3’ homology arm CRISPR oligonucleotides, respectively. Purple bar, probe used for Southern blotting. NLS, nuclear-localization signal; B, BglII recognition site. (B) Results of Southern blotting using the probe indicated in (A). (C) Evaluation of cytoplasmic and nuclear POU1F1 expression levels by immunoblotting. Anterior pituitary cells from four-week-old *Gh^wtGH1/wtGH1^* and *Gh^wtGH1 Δ3GH1^* mice were dissociated, separated into cytoplasmic (Cy) and nuclear (N) fractions, and subjected to immunoblotting analysis. GAPDH and LAMIN A/C were used as positive controls for Cy and N fractions, respectively. (D) Evaluation of wild-type GH, CREB3L1, and CREB3L2 expression in pituitary glands from 4-week-old *Gh^wtGH1/wtGH1^* and *Gh^wtGH1/Δ3GH1^* mice by immunofluorescence analysis.

---

**Supplementary table 1.**
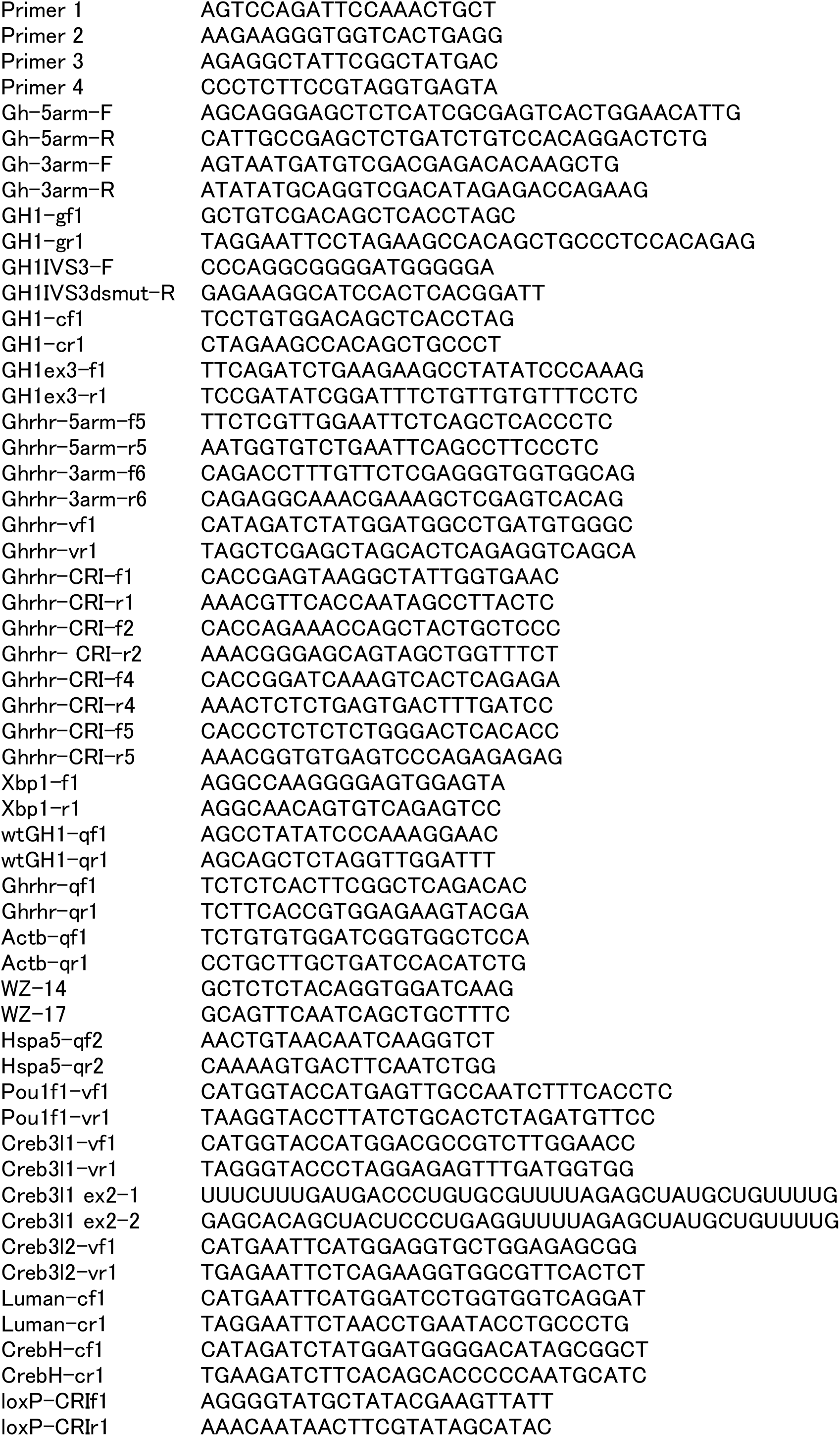
Sequences of primers used in this study.

